# Structural basis for membrane remodelling by the AP5:SPG11-SPG15 complex

**DOI:** 10.1101/2024.06.14.598999

**Authors:** Xinyi Mai, Yang Wang, Xi Wang, Ming Liu, Fei Teng, Zheng Liu, Ming-Yuan Su, Goran Stjepanovic

## Abstract

The human spastizin (spastic paraplegia 15, SPG15) and spatacsin (spastic paraplegia 11, SPG11) complex is involved in cargo sorting from late endosomes to the Golgi, and mutations in these two proteins are linked with hereditary autosomal recessive spastic paraplegia (HSP). SPG11-SPG15 can cooperate with evolutionarily ancient fifth adaptor protein complex (AP5). We employed cryo-electron microscopy and *in silico* predictions to investigate the structural assemblies of SPG11-SPG15 and AP5:SPG11-SPG15 complex. The W-shaped SPG11-SPG15 intertwined in a head-to-head fashion, and the N-terminal region of SPG11 is required for AP5 complex interaction and assembly. The AP5 complex is in a super open conformation. We employed *in vitro* lipid binding assays and cellular localization analysis to investigate AP5:SPG11-SPG15 membrane binding properties. Here we solve a major problem in understanding AP5:SPG11-SPG15 function in autophagic lysosome reformation (ALR), using a fully reconstituted system. We reveal that the AP5:SPG11-SPG15 complex binds PI3P molecules, can sense membrane curvature and drive membrane remodelling *in vitro*. These studies provide key insights into the structure and function of the spastic paraplegia AP5:SPG11-SPG15 complex, which is essential for the initiation of autolysosome tubulation.

## Introduction

Hereditary spastic paraplegia (HSP) is a group of inherited neurodegenerative and developmental diseases caused by the progressive degeneration of the corticospinal tracts, leading to weakness and spasticity of the lower limbs. More than 80 spastic paraplegia genes (SPGs) have been identified [1]. The most common form of autosomal-recessive HSP is caused by mutations in genes that encode two large ∼250 kDa proteins, SPG11 and SPG15 (also known as spatacsin and spastizin, respectively) [2–4]. To date, more than 300 mutations in the *SPG11* gene have been reported [5]. Most patients exhibit distinctive features of spastic paraplegia, cognitive impairment, early-onset parkinsonism, corpus callosum changes and polyneuropathy [3]. SPG11 is stably associated with SPG15 and the fifth adaptor protein complex (AP5), which is involved in late endosome-to-Golgi retrieval of cargo proteins [6, 7]. However, the precise function of AP5 and its associated proteins in this pathway is still largely unknown.

The AP5:SPG11-SPG15 complex is localized to late endosomes and lysosomes [8–11], and its loss results in lysosome abnormalities, including endolysosomal accumulation of material and enlargement, lack of free lysosomes available for fusion with autophagosomes and autophagosome accumulation [12–15]. Apart from HSP, autophagy and lysosomal defects are associated with a number of neurodegenerative conditions, including Parkinson’s disease, Huntington’s disease, and Alzheimer’s disease [16–19]. Thus, defects in this pathway may represent a new type of lysosome storage disease (LSD) with links to neurodegeneration. Mechanistically, SPG11 and SPG15 are believed to be essential for the initiation of lysosomal tubulation and autophagic lysosome reformation [7, 14]. SPG11 and SPG15 are predicted to have α-solenoid structures that are present in clathrin, α-COP, and β‘-COP [20]. Therefore, SPGs may be involved in AP5-mediated cargo protein sorting; however, this has not yet been experimentally demonstrated. SPG11 also has an N-terminal WD40 domain responsible for interaction with AP5, while SPG15 harbors a zinc-binding FYVE (present in Fab1, YOTB, Vac1, and EEA1) domain that can bind to phosphatidylinositol 3-phosphate (PI3P), allowing the AP5:SPG11-SPG15 complex to localize to the endolysosomal compartment [8, 21].

AP complexes are evolutionarily conserved heterotetramers that are involved in the sorting of cargo proteins into transport vesicles and trafficking between different membrane compartments. Five different AP complexes can be differentially recruited to the Golgi, endosomes, or plasma membrane. AP1, AP2 and AP3 interact with the clathrin coat, which is thought to be important for membrane deformation and to concentrate cargo proteins during the formation of transport vesicles. The clathrin coat lattice is composed of triskelions, an assembly of three clathrin heavy chains each bound to a single light chain. Functionally, AP1, AP3 and AP4 complexes are associated with sorting of proteins at the trans-Golgi network (TGN), AP2 facilitates clathrin-mediated endocytosis, and AP5 is involved in retrieval of Golgi proteins from late endosomes. The mechanisms of signal recognition and membrane recruitment have been well studied for other AP complexes, but not AP5. AP1-4 all recognize similar types of sorting signals on cargo proteins, including tyrosine-based (YXXØ) motifs via their μ subunits and dileucine-based ([DE]XXXL[LI]) signals that bind to the small σ subunit. All five AP complexes have similar subunit compositions, with two large subunits (β1–5 and α, γ, δ, ε, or ζ), a medium-sized subunit (μ1–5), and a small subunit (σ1–5). The core of the AP complex is defined by the assembly of the N-terminal helical solenoid (trunk) domains of the two larger subunits, the medium and small subunits. The AP core mediates sorting signal recognition and binding to phosphoinositides in target membranes. The C-termini of the two large subunits contain appendage domains that protrude outside of the core and are connected to the N-terminal trunk domains by flexible linkers or hinges. The AP5 β5 subunit lacks a hinge region, while the ζ subunit lacks both a hinge and an appendage domain. The μ subunit can be further subdivided into the N-terminal globular domain (N-μ) and the C-terminal extended domain (C-μ). Each subunit within the adaptor complex is responsible for a specific function: the μ and σ subunits can associate with sorting signals on cargo proteins; the hinge region of β1-3 interacts with clathrin, while the α/γ type subunits bind different accessory proteins through their appendage domains. Unlike AP1-AP3, the AP5 complex does not associate with clathrin but rather with SPG11 and SPG15, which are hypothesized to form a membrane coat-like subcomplex [22]. In this aspect, AP5 is similar to the AP4 complex, which lacks the clathrin-binding motif in the β4 subunit and is able to function without clathrin (reviewed in [23]). However, the vesicle budding mechanism of these clathrin-independent APs remains poorly understood.

The recruitment of APs to the targeted membrane involves coincident interactions with anionic lipids, small GTPases and sorting signals on the cytoplasmic domains of membrane cargo proteins [24]. For example, recruitment of AP2 to the plasma membrane requires phosphatidylinositol 4,5-bisphosphate (PIP2) and sorting signals, whereas recruitment of AP1 onto membranes requires small GTPase ADP-ribosylation factor 1 (Arf1), phosphatidylinositol 4-phosphate (PI4P) and sorting signals. AP1 interacts with GTP-bound Arf1 via its γ and β1 subunits and PI4P with the N-terminus of its γ subunit, while AP2 interacts with PIP2 via two sites located at the α and μ2 subunits.

In AP1 and AP2 closed states, the trunk domains from the two large subunits form a bowl-like structure. The C-μ2 domain is located at the center of the bowl in AP2, and this conformation occludes the tyrosine motif-binding pocket in the μ2 subunit by portions of the β2 subunit. The dileucine binding site in the σ2 subunit is similarly occluded by binding the N-terminus of the β2 subunit, and the clathrin binding box is buried in the center of the AP2 core. The binding of AP2 to PIP2 and AP1 to Arf1 induces a conformational change from a closed to an open conformation [25–27]. This conformational change exposes cargo and clathrin binding sites on µ/σ and β subunits, respectively, and thus enables AP complexes to nucleate assembly of clathrin lattice coats at appropriate membrane sites.

Furthermore, the μ domains of AP1 and AP2 are known to be phosphorylated at a conserved threonine residue, and this modification may influence binding to both cargo and clathrin (reviewed in [28]). Recruitment of AP5 onto intracellular membranes does not require Arf1; instead, it appears to depend on SPG11-SPG15 interaction with Rag GTPases and PI3P [21]. The molecular details of this interaction and whether AP5 can directly bind to phosphoinositides and/or small GTPases and undergo similar conformational changes are not known.

Due to a lack of structural information, how the AP5 complex binds to SPG11-SPG15, their membrane assembly, and the molecular mechanism remain elusive. Here, we present cryo-electron microscopy (cryo-EM) structures of the full-length SPG11-SPG15 complex alone and bound to the entire core of AP5 (full-length ζ and β5 complexed with the σ5 and μ5 subunits) and discuss how their organization influences AP5 conformation and promote membrane remodeling. We employed *in vitro* lipid binding assays and cellular localization analysis to investigate AP5:SPG11-SPG15 membrane binding and remodelling properties. Moreover, we demonstrated that disrupting the SPG11-AP5 interaction interface results in enlargement of the endolysosomal compartment. This work serves as a structural framework for understanding the assembly and dynamics of the spastic paraplegia AP5:SPG11-SPG15 complex and offers insights into mechanisms of autophagic lysosome reformation and retrograde trafficking.

## Results

### Structure of the SPG11-SPG15 complex

To structurally characterize the human SPG11-SPG15 assembly, we overexpressed and purified the complex from Expi293F cells using two-step affinity purification (Figure 1A-B). SPG11-SPG15 samples were analyzed by negative stain electron microscopy (Figure S1A). Two-dimensional (2D) class averages revealed a W-shaped particle approximately 30 nm in length. We then performed cryo-EM analysis of SPG11-SPG15 by collecting multiple datasets, as detailed in the Materials and Methods section. The combined data enabled us to obtain a map of the SPG11-SPG15 complex with an overall resolution of 4.02 Å and build the structure with clear sidechain identities in better resolved parts of the structure using the initial models generated by AlphaFold2 prediction (Figure S2). While our structure consists of a single SPG11-SPG15 heterodimer, 2D classification analysis from negative staining revealed a small percentage of particles that represent oligomeric assemblies consisting of two interacting protomers (Figure S3A-B). These observations support the hypothesis that the building units of SPG11-SPG15 complexes are heterodimers that can form different higher-order assemblies.

**Figure 1:**
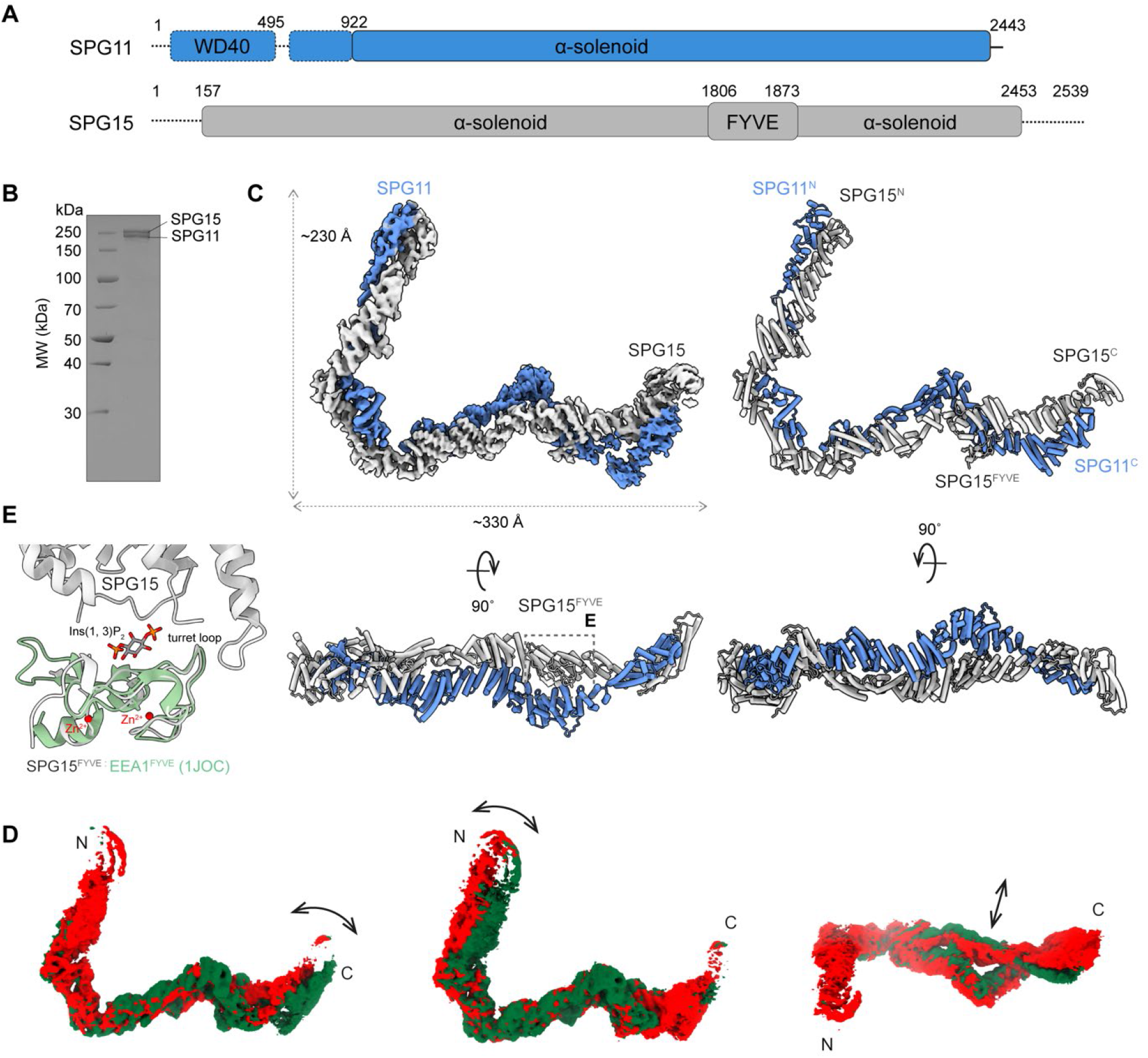
Architecture and dynamics of the human SPG11-SPG15 complex. **(A)** Domain organization of the human SPG11-SPG15 complex. Unresolved regions are indicated by dashed lines. **(B)** Coomassie blue-stained SDS-PAGE of purified SPG11-SPG15. MW, molecular weight. **(C)** Cryo-EM density map and the model of the human SPG11-SPG15 complex shown in different views. SPG15 and SPG11 are colored in light gray and royal blue, respectively. **(D)** Results of 3DVA with three variability component images of SPG11-SPG15. The red and green map indicate the movement between these two states. **(E)** The structural alignment of SPG15^FYVE^ with EEA1^FYVE^ (PDB: 1JOC). The FYVE domain contains Zn2+ binding sites and interacts with PI3P polar head group.

Structurally, SPG11 is predicted to include the N-terminal WD40 domain, followed by a larger α-solenoid, while SPG15 has the domain arrangement of α-solenoid-FYVE-α-solenoid (Figure 1A). Our experimental model shows that the α-solenoid domain of SPG15 binds in a parallel manner to the α-solenoid of SPG11, forming a curved structure that is ∼230 x 330 Å long (Figure 1C). The modeled part of the structure is composed of almost full-length SPG15 (residues 157-2453) and the C-terminal α-solenoid region of SPG11 (residues 922-2443) with an overlapping area of ∼4861.3 Å^2^. The α-solenoid region of SPG15 is composed of 107 α-helices. The SPG11 and SPG15 α-solenoids coil around each other through most of the core assembly to form tight dimer contacts. The N-terminal region of SPG11 that includes the predicted WD40 domain was not resolved in the cryo-EM reconstruction. The 2D class averages suggest that the WD40 domain is flexibly linked to the larger α-solenoid region of SPG11. This is reminiscent of the N-terminal WD40 domain of the clathrin heavy chain (CHC), which is connected to the distal α-helical solenoid fold by a flexible linker. Apart from the SPG11 N-terminal region, the SPG15 extreme C-terminus (residues 2454-2539) is poorly resolved in the electron density maps and does not participate in the heterodimer interface. We performed 3D variability analysis (3DVA) in cryoSPARC v4 to capture the conformational dynamics of the complex [29]. The results demonstrated that the two terminal arms undergo continuous movements relative to the middle part of the canonical W (Figure 1D).

The SPG11 α-solenoid makes a nearly 90° turn after residue P1906, and together with SPG15 creates a binding platform for the FYVE domain. The FYVE domain (residues 1806-1873) is linked to the SPG15 core assembly by a long linker (residues 1738-1805 and 1874-1908) that is not visible in electron density maps and is likely unstructured. Membrane binding of FYVE domains is driven by electrostatic interactions with PI3P headgroups and includes insertion of the exposed nonpolar ’turret loop’ residues into the hydrocarbon interior of the lipid bilayer [30–34]. In SPG15, the conserved PI3P polar head group binding site is sandwiched between the beta-strand core of the FYVE domain and α-solenoid, and the tip of the turret loop is inserted in the interface formed by SPG11 and SPG15 α-solenoids (Figure 1E). Therefore, when in solution, both membrane interaction sites appear to be in conformations that preclude PI3P engagement and membrane insertion. A conformation change may be required to bring the PI3P binding site in contact with the head group region of the lipid bilayer and release the hydrophobic residues in the turret loop (residues 1824-1832) from the SPG11-SPG15 dimer interface. The long linkers connecting the FYVE domain to the flanking α-solenoids would allow for FYVE domain reorientation when on the membrane surface.

### Structure of the AP5:SPG11-SPG15 complex

AP5 plays a role in regulating late endosomes to Golgi retrieval of various proteins, and together with the SPG11-SPG15 complex contributes to autophagic lysosome reformation. To date, the AP5 complex has not been structurally characterized, and the molecular mechanisms that regulate the interactions between these proteins as well as differences between AP5 and other adaptor protein complexes are not known. To determine the AP5 structure and visualize how SPG11-SPG15 associates with AP5, we reconstituted and purified the complex from Expi293F cells by coexpressing full-length mouse ζ and human β5 with full-length human σ5 and mouse μ5 (Figure S1B, S4). Initial attempts to reconstitute the AP5 complex resulted in purification of the ζ/σ5 and β5/µ5 subcomplexes, each pair forming a stable heterodimer that migrate as two separate peaks in size exclusion chromatography. Interaction with SPG11 residues 1-525, named the SPG11^WD40-hairpin,^ was necessary to assemble the two AP5 subcomplexes into a complete AP5 heterotetramer (Figure S5A). Pull-down analysis showed that SPG11 interacts primarily with the ζ/σ5 subcomplex (Figure S5B). The purified AP5:SPG11-SPG15 complex appeared as a stable heterohexamer judged by SDS‒PAGE (Figure S4A). Cryo-EM images were collected and processed as detailed in the Materials and Methods section, resulting in reconstruction of the AP5:SPG11-SPG15 complex at an average resolution of 3.30 Å (Figure S4). The 2D class averaging and 3D refinement procedures converged to the part of the complex containing AP5 bound to the previously unresolved N-terminal WD40 domain of SPG11. The extended SPG α-solenoids are highly variable and poorly resolved in the final reconstruction. This is consistent with 3D variability analysis showing a highly variable position of the N-terminal region of SPG11. We further fitted the AlphaFold2 models of the individual AP5 subunits and SPG11 WD40 domain into the map. The final reconstruction consists of two large subunits (ζ and β5) a medium-sized subunit (μ5), and a small subunit (σ5) of AP5 bound to the SPG11^WD40-hairpin^ domain (Figure S4D). The small σ5 subunit consists of a five-stranded β-sheet and several α-helices packed against it. σ5 binds to the ζ helical solenoid (trunk) domain, while the N-terminal domain of µ5 (N-µ5) binds to the central core of the β5 trunk domain (Figure S4G). The C-terminal domain of µ5 (C-µ5) has an elongated, all β-sheet structure and interacts exclusively with the β5 trunk domain. The highly conserved binding site for tyrosine-based endocytic motifs found in the μ subunit of AP1-4 is altered in µ5 (Figure 3I), indicating either loss of cargo binding functionality or change in specificity. As in the previously reported structures of AP1, AP2 and AP3, the appendage domain of β5 (residues 633-878) is joined to the rest of the protein via a flexible linker and was not resolved in the final reconstruction. Such flexibly connected appendage domains are thought to act as binding platforms for accessory proteins.

Next, we examined whether the appendage domain influences the assembly of AP5 with the SPG complex. We repeated the sample preparation using a truncated β5 construct (residues 1–632) lacking the appendage domain and collected additional single-particle cryo-EM data (Figure 2A-2C, Figure S6-S7). This AP5 structure was determined at a higher resolution (3.26 Å) compared to the complex containing full-length β5. This result can be explained by the reduced conformational heterogeneity of the AP5 core compared to that of the full-length complex. The most prominent difference is the poorly resolved density of the helical linker connecting the SPG11 WD40 domain and distal alpha helical solenoid, indicating higher flexibility of this region in the absence of the appendage domain (Figure 2). Pull-down assays indicated that the appendage domain mediates weak interaction between the β5 subunit and SPG11-SPG15 complex, providing possible explanation for differences in the relative mobility between the two complexes (Figure S5C). The SPG11 N-terminal region interacts with the C-terminal half of the ζ and β5 trunk domains and bridges the ζ/σ5 and β5/µ5 subcomplexes to form a stable heterotetramer (Figure 2C). The SPG11 N-terminal domain folds into a seven-bladed WD40 domain followed by two short α-helices (h1 and h2), forming a helical hairpin, i.e., the SPG11^WD40-hairpin^ (Figure 2D). This helical hairpin is part of a longer helical linker region that connects WD40 to the distal α-solenoid of SPG11 and was partially visualized in the reconstructions of the complex containing full-length β5 (Figure 2E). Each of the seven β-sheets is made up of four antiparallel strands denoted a to d (Figure 2D). The SPG11 WD40 structure is similar to the N-terminal propellers found in CHC, β’-COP and α-COP from the COPI coat as well as the Sec31 subunit of the COPII coat [35–37]. The most prominent difference occurs in blade V at the transition between the βc and βd strands. In clathrin, the β-hairpin is formed by the βc and βd strands oriented in an antiparallel direction and linked by a short loop. In SPG11, this loop is replaced by the ∼100-residue sequence that can adopt a partial α-helical structure (Figure 2D). In addition, the SPG11 β-propeller is arranged in a circular fashion, while the clathrin propeller has an elliptical cross section.

**Figure 2:**
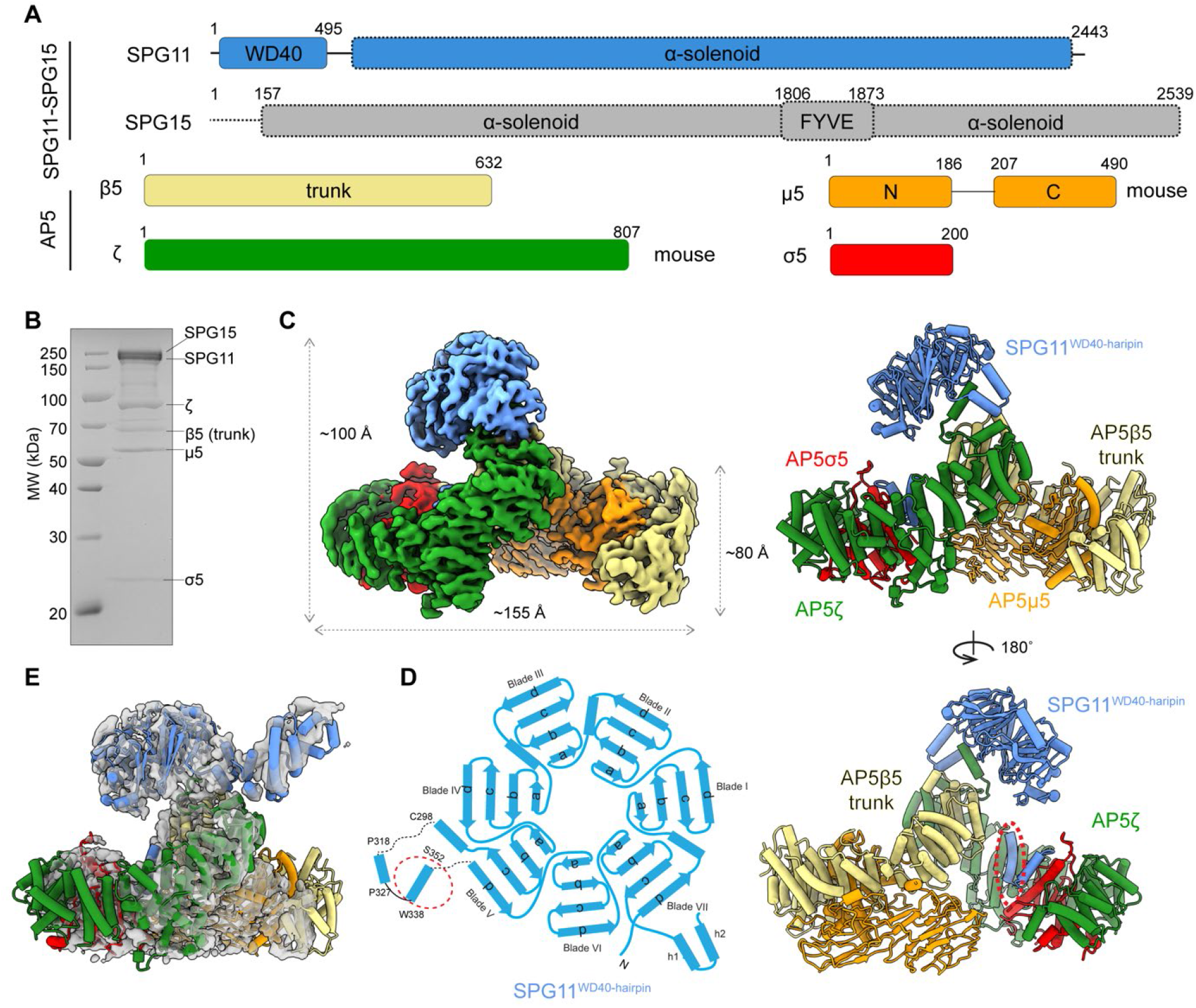
Cryo-EM structure of the AP5^βtrunk^:SPG11-SPG15 complex. **(A)** Domain organization of the AP5:SPG11-SPG15 complex. Unresolved regions are indicated by dashed lines. **(B)** Coomassie blue-stained SDS-PAGE of purified AP5^βtrunk^:SPG11-SPG15 complex. MW, molecular weight. **(C)** Cryo-EM density map and the model of the AP5^βtrunk^:SPG11-SPG15 complex shown in different views. SPG11^WD40-hairpin^ and AP5 are colored in cornflower blue, forest green (ζ), khaki (β5), red (σ5) and orange (μ5) respectively. The anchor motif of SPG11 is boxed and detailed residues are shown in (D). **(D)** Molecular model of SPG11^WD40-hairpin^. WD40-hairpin domain containing a seven-bladed WD40 domain, followed by two short α-helices (h1 and h2) forming a helical hairpin. The anchor motif of SPG11 is boxed in red circle. **(E)** The model of the AP5^FL^:SPG11-SPG15 complex fitted into the map showing extra density of the helical linker connecting the SPG11^WD40-hairpin^ and α helical solenoid.

**Figure 3:**
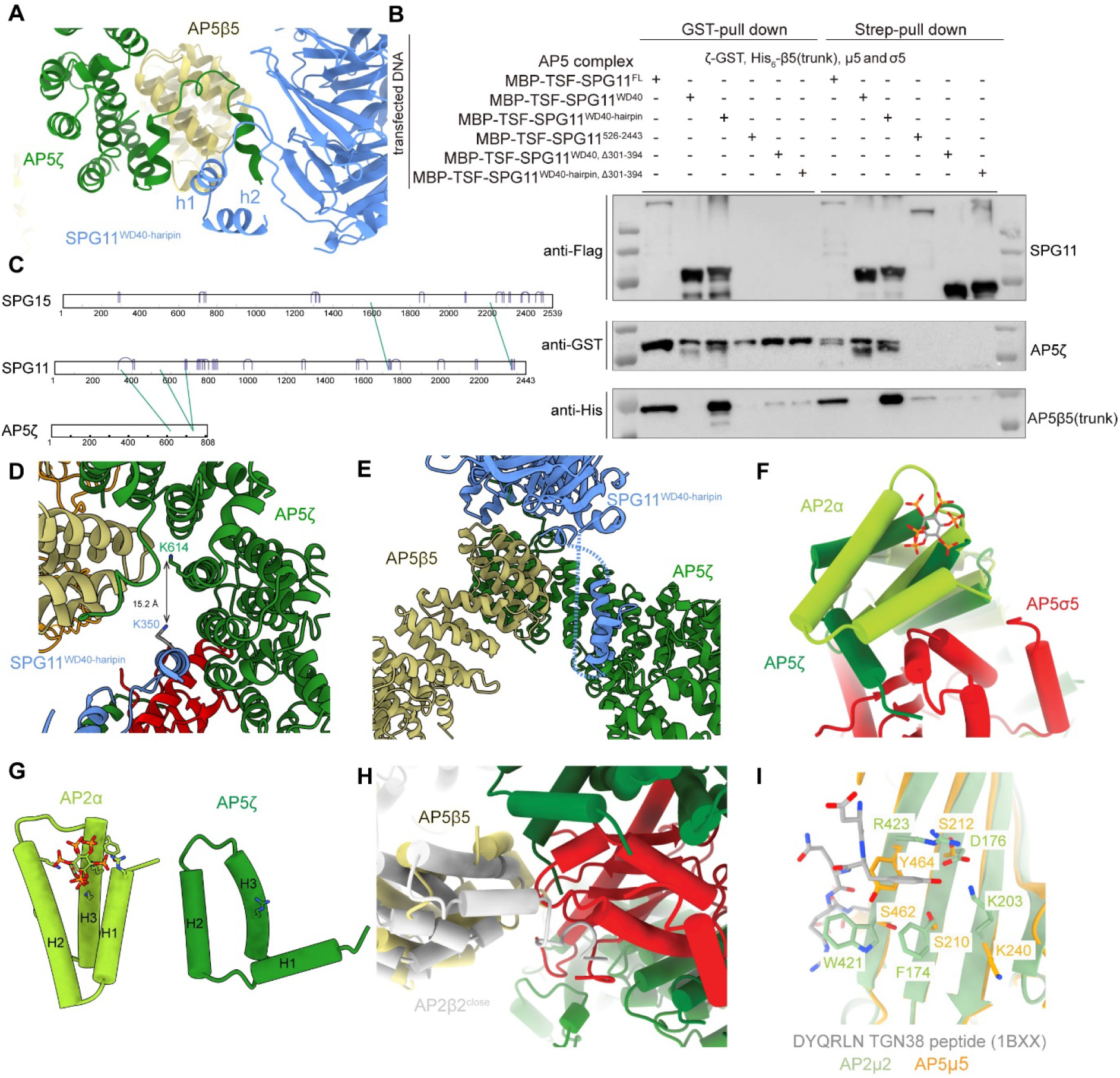
Interface between the SPG11^WD40-hairpin^, AP5β5 and AP5ζ. **(A)** Zoomed-in-view of the interface between the SPG11^WD40-hairpin^, AP5 β5 and AP5ζ. **(B)** Pull-down assay of full length or truncated MBP-TSF-SPG11 with AP5 complex. **(C)** Crosslinks between AP5:SPG11-SPG15 mapped onto its primary structure scheme. The detected intermolecular crosslinks include SPG11(K350) - AP5ζ(K614), SPG11(552) - AP5ζ(726), SPG11(678) - AP5ζ(726), SPG11(1740) - SPG15(1596), SPG11(2368) - SPG15(2216). **(D)** Crosslink mapped onto the SPG11-AP5 structure. The zoomed-in view of representative crosslink between SPG11^WD40-hairpin^(K350) and AP5ζ(K614) was shown. **(E)** Zoomed-in view of the interaction between the anchor motif of SPG11 and AP5 ζ. **(F)** Structure comparison of AP2α (PDB 2VGL) and AP5ζ, indicates the difference of AP2α PIP2 binding sites and corresponded AP5 ζ. **(G)** Side-by-side comparison between AP2α and AP5ζ. The sidechain of positive charged residues of AP5ζ and PIP2 binding sites of AP2α were shown. **(H)** Structure comparison of closed state AP2β2 and AP5β5. **(I)** Structural alignment between the μ domain of AP2 and AP5 bound to the tyrosine-based motif, DYQRLN TGN38 peptide (PDB:1BXX).

The interaction between the SPG11 propeller and AP5 complex is mediated by the helical hairpin that packs against the surface of the C-terminal α-helices of the ζ and β5 trunk domains (Figure 3A). The ζ subunit C-terminus (residues 781-788) folds into an α-helix that is positioned in a pocket created by the helical hairpin and the bottom surface of blade VII of SPG11. The C-terminal parts of the ζ and β5 trunk domains bind to each other with an overlapping area of ∼700 Å^2^. This is in sharp contrast to a much larger overlapping area of ∼1500 and ∼1600 Å^2^ in AP1 and AP3, respectively. The structural arrangement of SPG11 WD40 appears to stabilize the interaction between the ζ and β5 trunk domains by forming a more extensive interface. Deletion of the SPG11^WD40-hairpin^ (Δ1-525, i.e., 526-2443) completely abolished the interaction with the AP5 complex, as confirmed by coexpression and pull-down experiments. Conversely, using SPG11^WD40-hairpin^ as bait was sufficient to pull down both AP5 adaptins (Figure 3B). Next, we tested the importance of SPG11 h1 and h2 for this interaction. The shorter SPG11 construct, SPG11^WD40^ (residues 1-495), lacking both helices, was not able to pull down β5/µ5 but interestingly retained its interaction with ζ/σ5. Closer inspection of the model and map revealed disconnected electron density that binds to the ζ/σ5 subcomplex and could not be attributed to the main chain of the AP5 complex. In an attempt to identify and model this region, we performed chemical crosslinking combined with mass spectrometry (XL-MS). We used the MS-cleavable crosslinker DSBU (disuccinimidyl dibutyric urea), which connects primary amine groups of lysine residues within Cα-Cα distances up to ∼30 Å. XL-MS analysis revealed several intermolecular crosslinks, including SPG11(K350)-AP5ζ(K614), located in the extended region between the βc and βd strands from Blade V of SPG11 and the ζ trunk domain, respectively (Figure 3C-3D). Subsequent protein modeling resulted in the final reconstruction that satisfied all spatial restraints and revealed the WD40 domain α-helix (residues R336-S352) bound to the ζ/σ5 subcomplex (Figure 3E). We verified this interaction by deleting residues 301-394 from the SPG11^WD40-hairpin^ and SPG11^WD40^. Both SPG11 constructs failed to pull down ζ and β5 subunits, indicating that this “anchor” motif is necessary and sufficient for the interaction with the ζ/σ5 subcomplex and by extension required for the whole AP5 complex assembly (Figure 3B).

Previously, it was observed that SPG11- and SPG15-mutated fibroblasts show autophagic and lysosomal defects, including the accumulation of enlarged lysosomes and an increased number of autophagosomes [38, 39]. This prompted us to examine whether mutations in the SPG11 β-propeller that disrupt interactions with AP5 cause a similar enlargement of endolysosomal compartment. We carried out fluorescence imaging studies on cells stably expressing the late endosomal/lysosomal marker TMEM192. In HeLa cells depleted of SPG11 by siRNA, lysosomes were significantly enlarged compared with controls (Figure 4). The normal endolysosomal compartment size could be partially restored by re-expressing wild-type SPG11. All of the N-terminal constructs of SPG11 as well as the SPG11 (Δ1-525, i.e., 526-2443) mutant showed no significant recovery at similar levels of expression (Figure 4A-4B), suggesting that the endolysosomal dysfunction phenotype in SPG11-depleted cells depends on binding AP5. However, neither interaction with ζ nor assembly of the full AP5 complex by the β-propeller alone was sufficient to restore endolysosomal compartment size (Figure 4A-4B). These data suggest that not only the SPG11^WD40-hairpin^/AP5 module but also the extended helical solenoid of SPG11 is required to restore the phenotype, most likely by allowing for interaction with SPG15. Many of the SPG11 mutations causative of spastic paraplegia are predicted to result in an abnormally truncated protein [2, 3, 40, 41]. We mapped these mutations onto the SPG11-SPG15 model (Figure S8). This revealed presence of pathogenic mutations in the WD40 domain, helical linker and also the extended helical solenoid, further underlining the importance of both WD40 domain and the extended solenoid for SPG11 function.

**Figure 4:**
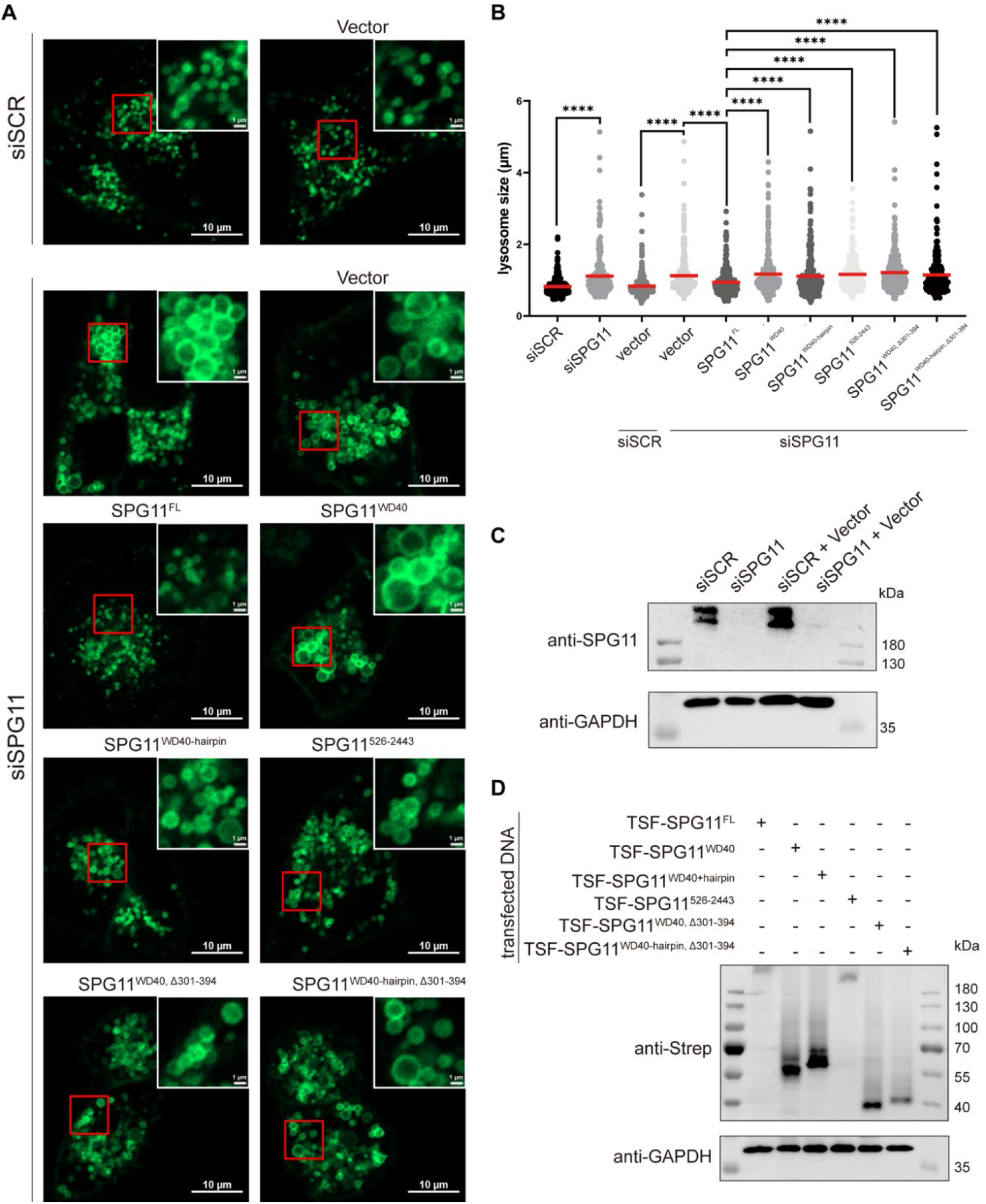
Disruption of the SPG11-AP5 interaction resulted in late endosomes/lysosomes enlargement. **(A)** After SPG11 downregulation, GFP-TMEM192-expressing HeLa cells were transfected with empty vector, wild-type SPG11, or truncated SPG11. Cells were imaged by confocal microscopy. Representative cells are shown. The inset shows an expanded view of the region in the red box. siSCR, scrambled control siRNA. **(B)** Quantification of the size of late endosomes/lysosomes in (A). (n = 100); Mean ± SEM are shown. Two-way ANOVA was used, ****: P < 0.0001. **(C-D)** Cells treated with siSCR or siSPG11 were transfected with empty vector, wild-type SPG11 or truncated SPG11, and cell lysates were immunoblotted. Molecular weight standards (in kDa) are shown.

### AP5 assumes a super-open conformation

Previously, diverse conformations in AP1 and AP2 were reported, including “closed”, “unlatched” “open” and “hyper-open” states. The cargo binding pockets in the small and medium subunits as well as the clathrin binding box in the β hinge domain are inaccessible in the closed structures. To expose these binding sites, AP complexes need to undergo a large conformational change to the open form. When the AP2 complex is localized to the appropriate membrane, the C-μ2 domain becomes dislodged from the complex core, exposing the tyrosine motif-binding pocket. The dileucine motif-binding pocket in the σ2 subunit and the clathrin binding box also become accessible, culminating in cargo protein and clathrin binding. Unlike previous conformations observed in other AP complexes, the AP5 core adopts a “super-open” conformation in solution, with two stretched arms as rectangles of approximately 155 Å (Figure S9). The C-μ5 subdomain is bound to the β5 trunk domain in a conformation that is similar to the active, open form of AP2 [25]. In the closed state of AP2, the dileucine motif-binding pocket on the σ2 subunit is occupied by the N-terminus of β2 [42]. We superimposed the N-terminal part of β2 from the closed AP2 complex (PDB 2VGL) with the structure of β5. A direct comparison between the two β subunits revealed that the β5 structure is shorter and missing the N-terminal sequence responsible for σ binding. Instead, the σ5 binding site is plugged by the nearby N-terminus of ζ (Figure 3F-3H). This structural arrangement, together with the unique interaction interface between the large adaptins (Figure 3A), appears to favor the open conformation of the AP5 complex. However, further particle classification and refinement of the complex containing full-length β5 revealed the existence of additional, more compact states of the open AP5 core, probably due to the conformational flexibility of β5 and ζ (Figure S4H). The super-open conformation results in co-planar arrangement of all theoretical cargo- and lipid-binding sites and likely represents the active form of AP5.

Structural alignment and comparison between different adaptor proteins revealed that AP5 is even more open than the “hyper-open” AP1 structure and the “stretched-open” conformation of AP3 (Figure S9). AP5 is an evolutionarily ancient complex that was the first to branch off from the rest of the AP lineage after their divergence from the COPI [43]. COPI contains the adaptor-like subcomplex that consists of four subunits with sequence and structural similarities to APs [44]. Structural comparison between AP5 and the membrane-bound adaptor-like subcomplex of COPI [35, 45] revealed that both complexes adopt a similar extended conformation that is even more open than previously observed for other active AP complexes (Figure S10). Furthermore, our data show that the SPG11-SPG15 complex is stably associated with AP5 in the absence of membranes (Figure S4). This is another similarity to the COPI coatomer, where the outer-coat and adaptor-like subcomplex form the cytosolic complex that is recruited to the membrane as an intact unit [46]. Taken together, SPG11 appears to play a key role in AP5 activation by stabilizing the β5-ζ solenoid-solenoid interface and allowing for the open complex formation.

### Membrane interactions of the AP5:SPG11-SPG15 complex

Adaptor proteins localize to appropriate membranes through coincident binding of cargo, phosphoinositides and the small GTPase Arf1. These interactions are mutually stabilizing, ensuring proper assembly of adaptor proteins and coat complexes [47]. It remains unclear whether AP5 interacts with phosphoinositides directly, as is the case with other APs [48–52], and how it contributes to the overall membrane targeting of the full complex. To investigate whether SPG11 depends on SPG15 and AP5 for its localization to late endosomes/lysosomes, we carried out fluorescence imaging on HeLa cells stably expressing TMEM192. SPG11 subcellular localization was analyzed in cells transiently transfected with SPG11-mCherry and SPG15. We found that both full-length SPG11 and SPG11 (526-2443) colocalized with TMEM192 when co-expressed with SPG15 (Figure 5A). In contrast, SPG11 localization to late endosomes/lysosomes was lost in cells that were not co-transfected with the SPG15 plasmid. These results indicate that SPG11 recruitment to the late endosome/lysosome membrane is AP5 independent but requires SPG15 for its localization. The initial localization of APs depends in part on the interaction with phosphoinositides found in relevant membranes and is mediated by the three N-terminal helices of α/γ-type subunits. To assess the influence of phosphoinositides on AP5 recruitment, we used an *in vitro* protein-lipid overlay assay. We evaluated the contribution of AP5 to phosphoinositide interactions using a minimal complex assembly consisting of all four subunits of AP5 and the SPG11 β-propeller domain (AP5:SPG11^WD40-hairpin^). A protein-lipid overlay assay indicated that AP5 does not interact with any of the tested phosphoinositides, including PI3P, PI4P and PIP2 (Figure 5B). A direct comparison between the α chain in AP2 (PDB 2VGL) and the ζ chain revealed a prominent structural difference at the N-terminal corner of ζ that corresponds to the α subunit PIP2 binding site (Figure 3F-3G). The N-terminus of ζ folds back and interacts with σ5, resulting in displacement of the first N-terminal helix. This structural rearrangement, combined with the reduced density of positively charged residues located within this region, explains why AP5 does not bind phosphoinositides.

**Figure 5:**
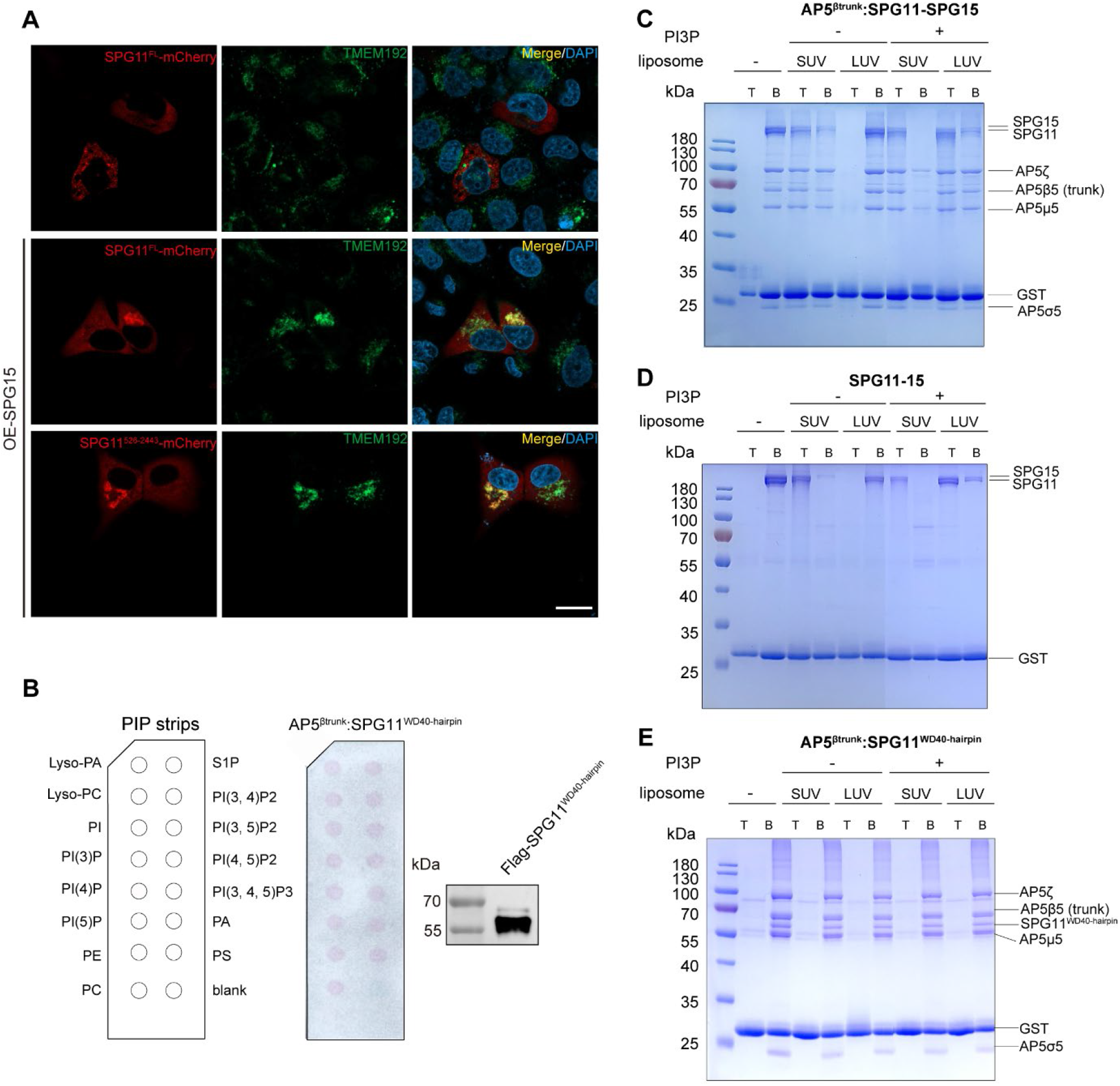
The membrane interactions of the AP5:SPG11-SPG15 complex. **(A)** HeLa cells that stably expressed GFP-TMEM192 were transfected with the indicated SPG11 constructs alone or further over-expressed with SPG15 and then processed for fluorescence detection of GFP and mCherry. OE, overexpression. Scale bar, 20 μm. **(B)** Recombinant Flag-tagged AP5^βtrunk^:SPG11^WD40-hairpin^ proteins were incubated with PIP strips. After binding and washing of the proteins to lipid spots, the strips were probed with antibodies to Flag and immunoblot reagents. **(C-E)** Liposome floatation experiments of AP5^Δtrunk^:SPG11-SPG15, SPG11-SPG15 and AP5^βtrunk^:SPG11^WD40-hairpin^ complexes were performed to test the interaction with LUVs and SUVs containing PI3P or not. GST protein was supplemented during TCA precipitation procedure to increase the protein recovery rate. T, top fraction; B, bottom fraction.

SPG11 and SPG15 were shown to be essential components in the initiation of autolysosome tubulation and autophagic lysosome reformation [7, 14, 53]. Thus, the AP5:SPG11-SPG15 complex could potentially localize at the membrane tubules due to a preference for highly curved membrane regions. To investigate this possibility, we performed liposome floatation experiments with both large unilamellar vesicles (LUVs; 100 nm) and highly curved small unilamellar vesicles (SUVs; ∼30 nm), either with or without 5% PI3P. In the experiments with LUVs, the top (lipid) fractions contained AP5^βtrunk^:SPG11-SPG15 and SPG11-SPG15 complexes only if PI3P was present in the membranes (Figure 5C-5D). Experiments with LUVs without PI3P showed that all proteins remained in the bottom fractions. As expected, the AP5^βtrunk^:SPG11^WD40-hairpin^ complex remained in the bottom fraction regardless of the presence of PI3P, LUVs or SUVs (Figure 5E). Therefore, the SPG11-SPG15 complex binds to membranes in a PI3P-dependent manner, consistent with the results of a previous study that demonstrated cytosolic localization of AP5, SPG11, and SPG15 upon cell treatment with the PI3-kinase inhibitor wortmannin [20]. In contrast, in the presence of SUV, AP5^βtrunk^:SPG11-SPG15 and SPG11-SPG15 complexes were mostly recovered from the top fractions in the presence or absence of PI3P, indicating that SPG11-SPG15 can sense lipid curvature and this may contribute to localization at the membrane tubules in addition to PI3P binding via the SPG15 FYVE domain.

### AP5:SPG11-SPG15 assemblies drive membrane remodeling *in vitro*

Previous work showed that the SPG11 and SPG15 together function in the initiation of auto-lysosomal tubulation and autophagic lysosome reformation [7, 14]. Our results suggest that AP5:SPG11-SPG15 complex could be the building unit of larger assemblies. To analyze the role of the individual subunits in membrane remodeling, we used giant unilamellar vesicles (GUVs) spiked with Rhod-PE, PI3P and DGS-NTA(Ni) lipids. GUVs were incubated with AP5^βtrunk^:SPG11-SPG15, SPG11-SPG15 or AP5^βtrunk^:SPG11^WD40-hairpin^ complexes, and monitored by confocal fluorescence microscopy (Figure 6A-6D). To monitor the localization of purified complexes directly, we tagged SPG11 with green fluorescent protein (GFP). AP5 interaction with cytoplasmic tails of potential cargo proteins was substituted with DGS-NTA lipids. DGS-NTA lipids allow binding of the AP5 to the membrane surface through the N-terminal 6His-tag in β5 which is located in the proximity of the predicted cargo binding site of μ5. The addition of the AP5^βtrunk^:SPG11-SPG15 complex led to membrane remodeling and produced protein-coated, outward membrane protrusions (Figure 6B). In contrast, the SPG11-SPG15 and AP5^βtrunk^:SPG11^WD40-hairpin^ complexes did not show any major membrane remodeling activity (Figure 6C-6D). These results show that AP5:SPG11-SPG15 complex is sufficient to introduce membrane reshaping *in vitro*, and that both AP5 and SPG11-SPG15 subcomplexes are required for this reaction. To test whether there is synergy between the AP5 complex, and SPG15-SPG11 in membrane remodelling, we prepared GUVs with and without PI3P or/and DGS-NTA. Membrane recruitment and remodelling reaction depended on PI3P on the GUV membrane (Figure 6B, S11B), in agreement with the PI3P–dependent recruitment of the AP5:SPG11-SPG15 complex *in vitro* (Figure 5C) and *in vivo* [20]. Furthermore, when we left out DGS-NTA from our GUV preparations, AP5^βtrunk^:SPG11-SPG15 complex was still able to localize to membranes, through interaction with PI3P, but failed to produce membrane protrusions (Figure S11C). The AP5:SPG11-SPG15 complex thus requires PI3P binding for membrane recruitment and remodelling while AP5 membrane interaction appear to primarily contribute to the membrane remodelling activity. These results are further corroborated with the finding that AP5:SPG11-SPG15 complex has the ability to sense membrane curvature and form larger assemblies which may contribute to membrane deformation.

**Figure 6:**
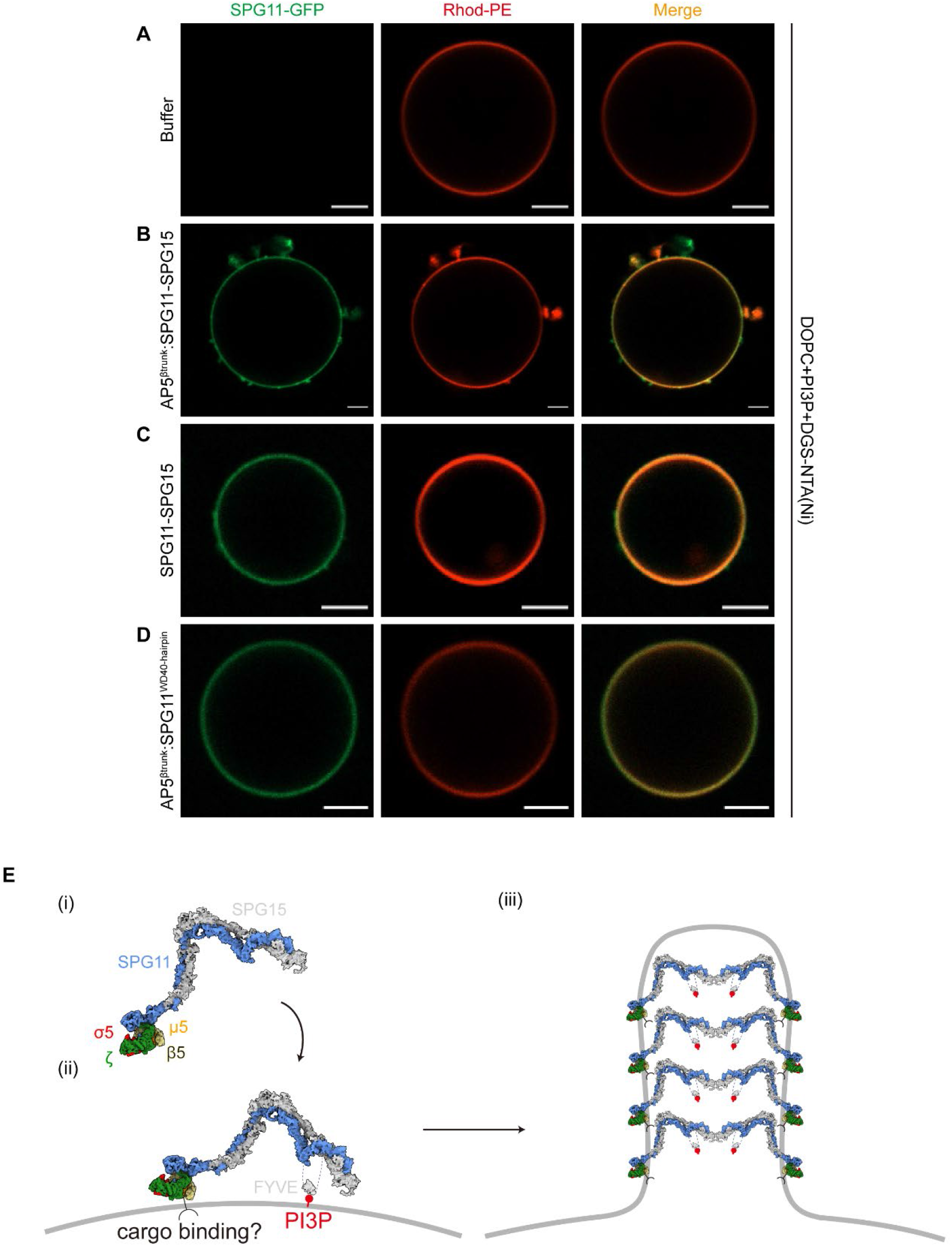
Effect of AP5^βtrunk^:SPG11-SPG15, SPG11-SPG15, and AP5^βtrunk^:SPG11^WD40-hairpin^ complexes on GUVs by fluorescence microscopy. **(A)** GUVs were shown by Rhod-PE fluorescence. **(B)** AP5^βtrunk^:SPG11-SPG15 complex caused membrane deformation as seen by several membrane protrusions around GUVs. **(C),(D)** SPG11-SPG15 or AP5^βtrunk^:SPG11^WD40-hairpin^ complex did not show remodeling effect on GUV membrane. Each experiment repeated independently 3 times with similar results. All scale bars are 5 μm. **(E)** Model of the AP5:SPG11-SPG15 complex interaction with PI3P and cargo. Assembly of the complex drives remodelling of endolysosomal membranes. Regulatory interactions with small GTPases are left out for simplicity. For further details, see the text.

## Discussion

Mutations in the autosomal recessive HSP gene products SPG11, SPG15 and the ζ subunit of AP5 result in lysosomal dysfunction, impaired membrane trafficking, autophagy defects with autophagosome accumulation and finally axon degeneration. However, the structure and molecular mechanism of these proteins have remained elusive. Here, we present the cryo-EM structure of the fifth adaptor protein complex, which shares a common evolutionary origin with COPI membrane trafficking complexes. AP5 forms a stable 1:1 complex with SPG11 and SPG15, two proteins with homology to clathrin heavy chain and COPI outer-coat subcomplexes. SPG11 and SPG15 form a membrane binding scaffold with the two proteins arranged in a head-to-head fashion that in turn interacts with AP5. The structure explains how SPG11 interacts with AP5 and SPG15 and provides mechanistic insights into the adaptor protein complex core. Our structure suggests that heterotetrametric AP5 complex assembly depends on interaction with the SPG11-SPG15 complex through the SPG11 WD40 domain. Here, we also show that deleting the SPG11 WD40 domain leads to endolysosomal compartment enlargement without affecting SPG subcellular localization. We also uncovered a previously unknown super-open conformation of AP5 with similarities to the adaptor-like subcomplex of COPI when bound to the membrane [35, 45]. The interaction between the SPG11 WD40 domain and AP5 is reminiscent of AP1 and AP2 interactions with the N-terminal β-propeller domain of the clathrin heavy chain, while the WD40 domains of α- and β‘-COP mediate cargo binding. In contrast to AP1 and AP2, which undergo a large conformational change from a closed to a membrane-bound open conformation that is permissive for clathrin binding, AP5 adopts an open conformation in solution with a coplanar arrangement of all potential membrane binding sites. More recent cryo-EM reconstructions of COPI and AP3 in their soluble forms have also revealed open conformations [52, 54]. Therefore, the evolutionary relationships between the AP5:SPG11-SPG15 complex, COPI and rest of the AP lineage appear to be reflected in the distinct structural characteristics.

Functionally, the AP5:SPG11-SPG15 complex was shown to play important roles in the initiation of autolysosomal membrane tubulation and autophagic lysosome reformation [6, 7, 14, 53]. A common feature of vesicular trafficking processes is donor membrane deformations induced by coat units, leading to budding and fission of membrane vesicles [55]. Similarly, autophagic lysosome reformation includes budding of tubular membrane structures. These tubules elongate along microtubules and finally undergo scission to produce nascent lysosomes [56]. Previous studies have suggested that AP5 acts as a backup system of retromer to promote the retrieval of transmembrane proteins from late endosomes to the trans-Golgi network [6]. The retromer complex and associated sorting nexins facilitate endosomal sorting of a variety of cargo molecules using tubular transport carriers [57–60]. Thus, both AP5:SPG11-SPG15 and the retromer complex are peripheral membrane proteins with important roles in the initiation of membrane tubulation and cargo sorting. The SPG11-SPG15 complex consists mainly of α-solenoids that assemble in an elongated and flexible domain to form prominent W-shaped arches. Similarly, arch-like assemblies are observed in the COPI adaptor-like and outer-coat subcomplexes as well as the retromer complex on the membrane [45, 57, 58]. The AP5 complex binds to the N-terminal edge of the arch, while the C-terminal part contains the FYVE domain. In this structural arrangement, SPG11 and SPG15 form an asymmetrical and flexible scaffold that can bind directly to the lipid bilayer via the FYVE domain. This bears similarities to the retromer complex, which also forms arch-shaped scaffolds and has a PX domain that specifically binds to PI3P-enriched membranes [57]. Furthermore, we showed that SPG11-SPG15 membrane localization does not depend on AP5. However, coincident recognition of additional factors such as small GTPases and/or potential membrane cargo proteins may contribute to AP5 membrane localization and function. Previous studies have demonstrated that SPG15 and SPG11 interact with Rab5A and Rab11 [61], as well as lysosomal resident Rag GTPases. When in isolation the SPG15 FYVE domain is recruited to the membranes *in vivo*, however in the context of the entire complex the FYVE domain exhibits activity only when the complex can interact with Rag GTPases [21]. In our structure the FYVE domain is found in a conformation that precludes membrane interaction when in solution. It is possible that interactions with regulatory proteins such as small GTPases induce a conformation change, releasing the FYVE domain which in turn may facilitate complex membrane recruitment and localisation to late endosomes and lysosomes. Furthermore, the conserved tyrosine motif binding site found in the μ subunits of AP1-4 is altered in AP5. Therefore, if AP5 is a cargo adaptor, it probably interacts with a different type of sorting signal, and their motifs remain to be determined. Local membrane deformations and budding induced by COPI assemblies are thought to be mediated by a combination of a curved scaffold and the insertion of Arf1 amphipathic helices. Similarly, in the case of retromer, membrane remodeling is driven by the combined action of the retromer core scaffold and membrane insertion of loops from the associated adaptor proteins. In all these cases, local membrane curvature is propagated over larger areas of membrane by coordinated oligomerization of a protein scaffold. Our results suggest that the AP5:SPG11-SPG15 scaffold has the potential to form larger assemblies, which may be functionally similar to other coat complexes on the membrane.

We propose a model to describe how the AP5:SPG11-SPG15 complex assembles on late endosome/lysosome membranes (Figure 6E). The model shows that the SPG11-SPG15 complex adapts an arch-like architecture that provides sufficient space for protein‒protein and protein-lipid interactions (i). AP5 is at the N-terminal tip of this arched structure, facing the membrane, and may in turn interact with membrane-cargo proteins. The C-terminal arm harbors the FYVE domain which anchors the complex to PI3P-enriched lipid bilayer (ii). Finally, the AP5:SPG11-SPG15 complex forms larger assemblies and deform membranes (iii). Our results show interdependence between AP5, PI3P, and SPG proteins for membrane binding and remodelling. The membrane remodelling activity must be controlled *in vivo*, and coincidence detection of membrane curvature, cargo and PI3P would allow SPGs to preferentially associate with tubular endolysosomal membranes. Further investigation of the organization of assembled AP5:SPG11-SPG15 complexes on auto-lysosomal tubules, and in particular its cross-talk with cargo, will be important for our understanding of SPG function in retrograde cargo sorting and ALR. In conclusion, our findings provide new insights into AP5:SPG11-SPG15 function and provide a structural context in which to test how SPG11 mutations cause autosomal recessive hereditary spastic paraplegia, and may open a new avenue for development of novel therapeutic strategies.

## Acknowledgments

We thank Dr. Chun-Yan Lim at Guangzhou National Laboratory for a gift of the TMEM192 cell line. The authors thank the cryo-EM (KEMC) and advanced mass spectrometry facility (KMS) of Kobilka Institute of Innovative Drug Discovery, the Chinese University of Hong Kong (Shenzhen). This work was supported by the Natural Science Foundation of Guangdong Province of China (to M.-Y.S., 2022A1515010856), the National Natural Science Foundation of China (31950410540 to G.S.), Foreign Young Talent Program from State Administration of Foreign Experts Affairs (QN2021032004L to G.S.), fund from Shenzhen-Hong Kong Cooperation Zone for Technology and Innovation (HZQB-KCZYB-2020056, to G.S.), CUHK-Shenzhen University Development Fund (to G.S.), and the Start-up funding from SUSTech (to M.-Y.S.).

## Contributions

X.M., M.-Y.S. and G.S. designed the experiments. X.M. and X.W. performed the experiments. X.M. and Y.W. collected the EM data. X.M., Y.W, and M.-Y.S. processed the EM data. F.T., M.L. and Z. L. contributed in the early stages of the project. M.-Y.S. and G.S. wrote the manuscript.

---

**Table S1:** Cryo-EM data collection, refinement and validation statistics.

|  | AP5<sup>βtrunk</sup>-SPG11-SPG15,<br>(EMDB-xxxx)<br>(PDB xxxx) | AP5<sup>FL</sup>-SPG11- SPG15<br>(EMDB-xxxx)<br>(PDB xxxx) | SPG11-SPG15 |
| --- | --- | --- | --- |
| **Data collection and processing** |  |  |  |
| Magnification | 105, 000 | 105,000 | 105,000 |
| Voltage (kV) | 300 | 300 | 300 |
| Electron exposure (e-/Å<sup>2</sup>) |  |  |  |
| Defocus range (μm) | -1.2 to -1.8 | -1.2 to -1.8 | -1.2 to -1.8 |
| Pixel size (Å) | 0.85 | 0.85 | 0.85 |
| Symmetry imposed | C1 | C1 | C1 |
| Initial particle images (no.) | ~2.6 millions | ~ 7 millions | ~6 millions |
| Final particle images (no.) | 399,340 | 586,806 | 234,969 |
| Map resolution (Å)<br>FSC threshold | 3.26 | 3.3 | 4.02 |
| **Refinement** |  |  |  |
| Initial model used (PDB code) | AlphaFold2 model | AlphaFold2 model | AlphaFold2 model |
| Model composition |  |  |  |
| Nonhydrogen atoms | 14200 | 14300 | 16481 |
| Protein residues | 2254 | 2274 | 3321 |

| Ligands | 0 | 0 | 0 |
| --- | --- | --- | --- |
| *B* factors (Å<sup>2</sup>)<br>Protein<br>(min/max/mean) | 52.13/360.99/148.66 | 81.15/560.47/175.52 | 68.84/381.80/172.87 |
| R.m.s. deviations<br>Bond lengths (Å)<br>Bond angles (°) | 0.003<br>0.607 | 0.004<br>0.678 | 0.013<br>0.912 |
| Validation<br>MolProbity score<br>Clashscore<br>Poor rotamers (%) | 1.78<br>8.53<br>0.21 | 1.96<br>11.97<br>0.52 | 1.87<br>8.82<br>0.00 |
| Ramachandran plot<br>Favored (%)<br>Allowed (%)<br>Disallowed (%) | 95.47<br>4.44<br>0.09 | 94.65<br>5.17<br>0.18 | 94.13<br>5.81<br>0.06 |
| CC (mask)<br>CC (box) | 0.71<br>0.83 | 0.63<br>0.81 | 0.69<br>0.75 |

## Materials and Methods

### Cloning and mutagenesis

The genes for mouse AP5ζ, mouse AP5μ5, human SPG11 and human SPG15 were codon optimized and synthesized. The genes encoding human AP5σ5 and human AP5β5 were amplified from human cDNA by PCR and subcloned into pCAG vectors with different tags using ClaI, KpnI and XhoI restriction sites. DNAs coding for the four subunits of the AP5 complex were as follows: mouse AP5ζ-GST (residue 1-807); human His_6_-AP5β5 (residues 1-632 for trunk domain or 1-878 for full length); mouse μ5 (residues 1-490); and human σ5 (residues 1-200). All four DNAs were subcloned into pCAG vectors separately. The TEV cleavage site was introduced between the GST tag and mouse AP5ζ. For the AP5:SPG11-SPG15 complex, human SPG11 was constructed with an N-terminal Twin-Strep-Flag tag, and human SPG15 was cloned with an N-terminal MBP tag followed by a GGGGSGGGGS linker and TEV cutting site. For the SPG11-SPG15 complex, human SPG11 was an N-terminal Twin-Strep-Flag tag, and human SPG15 was cloned with an N-terminal GST tag followed by a GGGGSGGGGS linker and TEV cutting site. For SPG11^FL^, SPG11^WD40^, SPG11^WD40-hairpin^, SPG11^526-2443^, SPG11^WD40, Δ301-394^, SPG11^WD40-hairpin, Δ301-394^ used in the pull-down experiment, they were cloned into the pCAG vector as an N-terminal MBP tag followed by a Twin-Strep-Flag tag. For AP5 subcomplex purification, AP5ζ and AP5β5 (1-632) were cloned as an N-terminal GST tag and a Twin-Strep tag in the C-terminus, respectively. For the AP5 subcomplex used in the pull-down experiment, mouse AP5ζ was cloned with a C-terminal GST tag, while human AP5β5 (full length or trunk domain) was constructed with an N-terminal GST tag.

### Protein expression and purification

For the AP5^FL^:SPG11-SPG15 and AP5^βtrunk^:SPG11-SPG15 complexes, N-terminal Twin-Strep-Flag-tagged human SPG11, N-terminal MBP-tagged human SPG15, and four subunits of the AP5 complex were coexpressed in Expi293F cells. Expi293F cells were grown in Union-293 medium and used for protein expression, with 1 mg of total DNA and 4 mg of PEI (Polysciences, 24765) per 1 liter of cells at a density of 1.5-2 x 10^6^ cells/ml. Cells were harvested after 3 days. The cell pellet was resuspended in lysis buffer (20 mM HEPES-NaOH pH 7.5, 150 mM NaCl, 2 mM MgCl_2_, 1 mM TCEP, 10% (v/v) glycerol) with protease inhibitors (1 mM PMSF, 0.15 μM aprotinin, 10 μM leupeptin, 1 μM pepstatin) and disrupted by sonication. After centrifugation at 18,000 rpm for 45 min, the cell supernatant was incubated with GST beads at 4 °C for 2 h. The beads were washed with wash buffer (20 mM HEPES-NaOH pH 7.5, 150 mM NaCl, 2 mM MgCl_2_, 2.5 mM DTT). Bound proteins were eluted with gel filtration buffer (20 mM HEPES-NaOH pH 7.5, 150 mM NaCl, 2 mM MgCl_2_) containing 50 mM reduced glutathione. After TEV digestion overnight at 4 °C, the elution was then subjected to Strep-Tactin Sepharose resin, washed with gel filtration buffer and eluted with gel filtration buffer supplemented with 10 mM desthiobiotin.

For the SPG11-SPG15 complex, N-terminal Twin-Strep-Flag-tagged human SPG11 and N-terminal GST-tagged human SPG15 were cotransfected into Expi293F cells as mentioned before. After 12 h, the cells were supplemented with 10 mM sodium butyrate. The transfected cells were cultured for an additional 48 h before harvesting. The harvested cells were washed with 1X cold PBS. The SPG11-SPG15 complex was purified using the same protocol as described above.

For AP5 subcomplexes, N-terminal GST tag and C-terminal Twin-Strep tag mouse AP5ζ and human AP5σ5 or N-terminal GST tag and C-terminal Twin-Strep tag human AP5β5 and mouse AP5μ5 were coexpressed in Expi293F cells with PEI. The cell pellet was lysed in lysis buffer supplemented with 1% (v/v) Triton X-100 for 30 minutes at 4 °C. Then, the AP-5 subcomplexes were purified as described before. After two-step purification, the protein elution was further purified by Superdex 200 Increase 10/300 GL (GE Healthcare).

For the AP5^βtrunk^:SPG11^WD40-hairpin^ complex, N-terminal Twin-Strep-Flag tag human SPG11 and four AP-5 subunits were cotransfected into Expi293F cells. The purification was the same as AP-5 subcomplex purification.

### Negative stain EM imaging

Four microliters of purified protein (0.03 mg/ml) was loaded onto a freshly glow-discharged carbon-coated 300 mesh grid (EMCN, BZ11023a) for 1 minute. The grids were subsequently blotted with filter paper and negatively stained with 2% (w/v) uranyl acetate for 40 seconds. Redundant liquid was absorbed using filter paper. The sample was imaged using a Talos 120C transmission electron microscope (Thermo Fisher) performed at 120 kV in low-dose mode and imaged with a Ceta CMOS camera (Thermo Fisher). Data were collected at a nominal magnification of 73000X, corresponding to a pixel size of 1.96 Å.

### Cryo-EM grid preparation and data acquisition

Purified AP5^FL^:SPG11-SPG15 and AP5^βtrunk^:SPG11-SPG15 complexes were prepared for cryo-EM data acquisition. For the cryo-EM grid, 4 μl of 1.5 mg/ml purified AP5^βtrunk^:SG11-SPG15 was applied to freshly glow-discharged Ultrafoil 1.2/1.3 300 mesh grids and plunged into liquid ethane using an FEI Vitrobot Mark IV set to 100% humidity, 4 °C, blot force 0, and blot time 3 sec after waiting for 5 sec. A total of 5724 movies were collected on a Titan Krios electron microscope operating at 300 kV equipped with a Gantan K3 camera at a defocus of -1.2 μm to -1.8 μm in counting mode, corresponding to a pixel size of 0.85 Å. Automated image acquisition was performed using SerialEM with a 3x3 image shift pattern. The movies consist of 50 frames, with a total dose of 61.86 e^-^/Å^2^, a total exposure time of 2.5 sec and a dose rate of 17.88 e^-^/pixel/sec. Imaging parameters for the dataset are summarized in Table S1.

For the AP5^FL^:SPG11-SPG15 complex, 4 μl of 0.9 mg/ml purified protein was deposited onto freshly glow-discharged Ultrafoil 1.2/1.3 300 mesh grids and plunged into liquid ethane using the same protocol as described above. A total of 10869 movies were collected on a Titan Krios electron microscope operating at 300 kV equipped with a Gantan K3 camera at a defocus of -1.2 μm to -1.8 μm in counting mode, corresponding to a pixel size of 0.85 Å. Automated image acquisition was performed using SerialEM with a 3x3 image shift pattern. The movies consist of 50 frames, with a total dose of 50.33 e^-^/ Å^2^, a total exposure time of 2 sec and a dose rate of 18.18 e^-^/pixel/sec. Imaging parameters for the dataset are summarized in Table S1.

The purified SPG11-SPG15 complex was supplemented with 0.02% β-octylglucoside immediately before freezing. Four microliters of 0.3 mg/ml protein sample supplemented with 0.02% β-octylglucoside was applied to freshly glow-discharged Ultrafoil 1.2/1.3 300 mesh grids and plunged into liquid ethane using the same protocol as the SPG11-SPG15:AP-5 complex. Four sets of movies were collected on a Titan Krios electron microscope operating at 300 kV equipped with a Gantan K3 camera at a defocus of -1.0 μm to -1.6 μm in counting mode, corresponding to a pixel size of 0.85 Å. Four datasets consist of 6660, 6849, 10746 or 9699 movies. Automated image acquisition was performed using SerialEM with a 3x3 image shift pattern. The movies consist of 50 frames, with total doses of 61.93 e^-^/ Å^2^, 57.58 e^-^/ Å^2^, 62.08 e^-^/ Å^2^, or 55.29 e^-^/ Å^2^, respectively. Imaging parameters for the datasets are summarized in Table S1.

### Crosslinking mass spectrometry (XL-MS)

Purified AP5^βtrunk^:SPG11-SPG15 was incubated with DSBU crosslinker (Thermo Fisher, A35459) to a final concentration of 200 μM for 40 min at 25 °C and quenched with 10 mM Tris-HCl for 10 min at 25 °C. The proteins were denatured with 10 mM DTT and 8 M urea for 60 min and then alkylated with 50 mM chloroacetamide (CAA) for 30 min in the dark. Then, the protein was digested by trypsin (trypsin: protein = 1:20, w/w) overnight at 37 °C. Tryptic peptides were desalted using Pierce peptide desalting spin columns (Thermo Fisher, 89852) and loaded (1000 ng in 0.1% formic acid) onto a nanotrap column (75 μm i.d. x 2 cm precolumn, packed with Acclaim PepMap100 C18, 3 μm, 100 Å; Thermo Fisher) using an EASY-nLC 1200 system (Thermo Fisher). Subsequent separation was performed on the analytical column (50 μm x 15 cm, Acclaim RSLC C18, 2 μm, 100 Å; Thermo Fisher) using a first gradient ranging from 2 to 8% of buffer B (80% acetonitrile and 0.1% formic acid) for 5 minutes followed by a second gradient ranging from 8 to 43% of buffer B for 80 minutes and a third gradient ranging from 43 to 50% of buffer B for 5 minutes at an overall flow rate of 300 nl/min. Nano-electrospray ionization was employed to ionize the samples. Ionized peptides were analyzed with an Orbitrap Eclipse mass spectrometer (Thermo Fisher) using the sceHCD-MS2 fragmentation method. Data-dependent analysis was carried out, using a resolution of 60,000 (AGC 4e5, max injection time 50 ms) for the full mass spectrum in the orbitrap. MS spectra were collected over a m/z range of 350-1600. MS/MS scans were recorded in the orbitrap at resolution of 30,000 (AGC 1e5, max injection time 120 ms, isolation width 1.6 m/z). Unknown, singly and doubly charged ions were removed from fragmentation. Selected precursors were fragmented using stepped higher-energy collision-induced dissociation (HCD) at normalized collision energies of 27%, 30%, and 33% NCE. Cross-linked peptides were analyzed and identified with Proteome Discoverer 2.5 and XlinkX software 2.5 (Thermo Fisher). The following settings were applied: two missed cleavage sites for trypsin per peptide were allowed; cysteine carbamidomethylation as fixed modification and methionine oxidation as dynamic modification. Searches were performed against a database including the sequences of AP5 complex, SPG11 and SPG15 and common contaminant proteins (CRAPome/Strep tag AP). Search results were filtered based on precursor tolerance (±10 ppm) and fragment tolerance (±20 ppm). The FDR threshold was set to 1% at the crosslink and CSM level.

### Cryo-EM data processing

For all the datasets, the movies were first aligned using MotionCor2 implementation in Relion 3 to correct the specimen movement and then imported into cryoSPARC v4 [62–64]. CTF (contrast transfer function) fitting and estimation were performed by patch ctf estimation. Four datasets were collected for the full-length SPG11-SPG15 complex. After particle picking from the initial subsets of 500 micrographs with a blob picker in the first dataset and applying to iterative cycles of 2D classifications, the good classes were chosen as templates for template-based particle picking, which generated approximately 6 million particles that were further subjected to 2D classification. The cleaned 609,170 particles were subjected to *ab initio* reconstruction to generate 3 classes. The best class was selected for nonuniform refinement, followed by global CTF refinement. The final map obtained from 234,969 particles was estimated at 4.02 Å. To improve the quality of the map, particle subtraction and local refinement were applied, resulting in 4.69 Å and 4.12 Å maps.

For the SPG11-SPG15:AP5 complex, 10,869 movies were collected. After particle picking from the initial subsets of 1500 micrographs with a blob picker and applying to iterative cycles of 2D classifications, the good classes were chosen as templates for template-based particle picking, which generated approximately 7 million particles that were further subjected to 2D classification. The cleaned 586,806 particles were subjected to *ab initio* reconstruction and nonuniform refinement. After global and local CTF refinement processing, the final reconstruction was estimated at 3.30 Å.

For the SPG11-SPG15:AP5^βtrunk^ complex, 5,724 movies were collected and processed with a similar procedure with additional heterogeneous refinement. The final map obtained from 399,340 particles was estimated at 3.26 Å.

The map was postprocessed using a deepEMhancer with tight target modes for model building [65]. All reported resolutions were based on the gold standard FSC 0.143 criterion.

### Atomic model building and refinement

The in silico models for AP5 components and SPG11-SPG15 were generated from AlphaFold2 structure predictions [66]. The backbones of the SPG11, SPG15 and AP5 subunits were docked separately into the density map using UCSF Chimera and further rebuilt by hand in Coot [67, 68]. Atomic coordinates were refined by iteratively performing Phenix real-space refinement and manual inspection and correction of the refined coordinates in Coot [69, 70]. To avoid overfitting, the map weight was set to 1, and secondary structure restraints were applied during automated real-space refinement. Model quality was assessed using MolProbity and map-vs-model FSC [71]. Figures were prepared using UCSF Chimera version 1.15 or ChimeraX [72]. The cryo-EM density map was deposited in the Electron Microscopy Data Bank under accession codes XXXXX and XXXXX, and the coordinates were deposited in the Protein Data Bank under accession numbers XXXXX and XXXXX.

### *In vitro* pull-down assay and western blot experiment

Different TSF-tagged SPG11 truncations and four subunits of the AP5 complex were transfected into Expi293F cells. The cells were harvested after 3 days of transfection and washed with 1X PBS. The cells were lysed as described previously. After centrifugation, the supernatant was divided into halves and incubated with GST or Strep-tactin Sepharose resin for 2 h. The beads were then washed with wash buffer and eluted with gel filtration buffer with 50 mM reduced glutathione or 10 mM desthiobiotin. The eluted samples were detected by western blotting with antibodies against Flag tag (CWBIO, CW0287), GST tag (Beyotime, AF5063), and His tag (CWBIO, CW0286). The experiment was repeated three times independently.

To test which subcomplex primarily interacts with SPG11, the GST-tagged ζ/σ5 subcomplex or GST-tagged β5/μ5 subcomplex was cotransfected with TSF-tagged SPG11 and MBP-tagged SPG15 into Expi293F cells. The pull-down assay and western blot were performed using the same protocol. The experiment was repeated three times independently.

### Liposome preparation

Liposomes were prepared by mixing lipid stocks dissolved in chloroform in glass test tubes in the following molar ratio: 75% DOPC:25% DOPE or 75% DOPC:20% DOPE:5% phosphatidylinositol-3-phosphate (PI3P). Lipid mixtures were evaporated to dryness under a nitrogen stream and then dried further under vacuum overnight. Dried lipid films were rehydrated with liposome buffer (20 mM HEPES-NaOH pH 7.5, 150 mM NaCl). Then, the mixture was vortexed vigorously to generate crude liposomes. Liposomes were subjected to 10 freeze‒thaw cycles to eliminate multilamellar structures. The final concentration of lipid was 2 mM. Small unilamellar vesicles (SUVs) were prepared by tip-probe sonication with 100 W, 3 s on, 5 s off for 10 minutes. Large unilamellar vesicles (LUVs) were prepared with a mini-extruder (Avanti Polar Lipids, 610000). Prepared unilamellar vesicles were subjected to the mini-extruder with a 0.1 μm polycarbonate membrane for 21 strokes.

### Liposome flotation assay

Proteins (0.5 μM) and liposomes (2 mM) were incubated in gel filtration buffer on ice for 30 minutes. The suspension was adjusted to 40% (w/v) OptiPrep (Sigma, D1556) by adding 48% (w/v) OptiPrep in the same buffer to a total volume of 300 μl, and the sample was transferred to an open-top thickwall polycarbonate tube (Beckman Coulter, 355635). A layer of 2.1 ml of 30% (w/v) OptiPrep was placed on top of each sample, and a layer of 50 μl liposome buffer supplemented with 0.5 mM TCEP was then placed on top. Reactions were centrifuged at 50,000 rpm for 3.5 hours in a Beckman SW60Ti rotor. The top and bottom fractions were collected and then TCA-precipitated and analyzed by SDS/PAGE.

### Protein-lipid overlay assay

The PIP strips (Echelon Biosciences, P-6001) were incubated with the AP5:SPG11^WD40-hairpin^ complex according to the manufacturer’s instructions. Briefly, the strips were first blocked with 3% fatty acid-free bovine serum albumin (BSA) in Tris-buffered saline containing 0.1% Tween 20 (TBS-T) for 1 hour. Then, 0.5 μg/ml purified protein in TBS-T/3% BSA was incubated with the PIP strips for 2 hours with gentle agitation. Bound proteins were detected with mouse anti-Flag (CWBIO, CW0287, diluted in 1:2000) primary antibody and an anti-mouse IgG-HRP secondary antibody (CWBIO, CW0102, diluted in 1:2000). The PIP strips were washed 30 minutes with TBS-T between each step. The whole procedure was performed at room temperature.

### Fluorescence microscopy experiments using giant unilamellar vesicles (GUV)

GUVs were prepared by the electroformation technique using Vesicle Prep Pro (Nanion). The lipid mixtures (80% DOPC:10% cholesterol:5% PI3P:5% DGS-NTA(Ni), 85% DOPC:10% cholesterol:5% PI3P:85% DOPC:10% cholesterol:5% DGS-NTA(Ni), or 90% DOPC:10% cholesterol) dissolved in chloroform at concentration of 5 mM was mixed with 0.1% of 18:1 Liss Rhod-PE, and then the lipid solution was spread onto the indium tin oxide (ITO) surface. The lipid was dried under vacuum for 1 hour to form a uniform lipid film. Vesicles were grown in a sucrose solution with the same osmolarity as the experimental buffer whereas an electric field (3 V, 5 Hz frequency) was applied for 2 hours at 37°C.

To study the membrane deformation effects of AP5^βtrunk^:SPG11-SPG15, SPG11-SPG15, and AP5^βtrunk^:SPG11^WD40-hairpin^ complex, proteins were added at a final concentration of 1μM to GUVs, respectively, with SPG11 or SPG11^WD40-hairpin^ tagged with GFP. GUVs were incubated with protein for 15 minutes at room temperature and imaged under a Zeiss LSM 900 confocal microscope with a Plan-Apochromat 40×/0.95 objective. Three independent experiments from three different GUV preparations were performed.

### Cell Culture and Transfection

HeLa cells stably expressing Flag-GFP-TMEM192 were grown in Roswell Park Memorial Institute (RPMI) medium (Eallbio, 03.4007C) supplemented with 10% v/v fetal bovine serum (FBS) and 1 μg/ml puromycin at 37 °C and 5% CO_2_. The cell lines were maintained by passaging the cells with trypsin-EDTA solution (Gibco, 25200056) after the density reached 75%-90% in a 60 mm dish (Corning, 430166). Transient transfections were performed using X-tremeGENE HP DNA Transfection Reagent (Roche, 06366236001) according to the manufacturer’s instructions.

### RNA interference

Knockdown was performed using the following On-Target Plus siRNA reagents from Dharmacon or a nontargeting SMARTpool siRNA as a control. The siRNAs were as follows: SPG11 (FLJ21439), J-017138-05, J-017138-06, J-017138-07, and J-017138-08. Knockdown was performed with Lipofectamine RNAiMAX (Thermo Fisher, 13778075) according to the manufacturer’s instructions using a final concentration of 25 nM siRNA in the culture medium.

### Western blotting

HeLa cells were plated into 12-well plate, followed by siRNA transfection using Lipofectamine RNAiMAX the next day, following the manufacturer’s instructions. Twenty-four hours after siRNA transfection, the cells were replated into a 12-well plate for transient plasmid transfections the next day. Cells were harvested 48 hours after plasmid transfection. After washing with PBS, cells were lysed in cell lysis buffer (50 mM Tris-HCl, pH 7.4, 150 mM NaCl, 0.5 mM EDTA, 0.5 mM MgCl_2_, 1% NP-40) supplemented with protease inhibitors (1 mM PMSF, 0.15 μM aprotinin, 10 μM leupeptin, 1 μM pepstatin) for 15 minutes on ice. Finally, insoluble material was removed by centrifugation (15000 rpm, 10 minutes), and total lysates were recovered. The protein concentration of the lysates was determined using a BCA protein assay (Sangon, C503021). Samples were loaded at equal amounts for SDS‒PAGE. PageRuler Plus Prestained Protein Ladder (Thermo Fisher, 26616) was used to estimate the molecular sizes of bands. Proteins were transferred to 0.22 μm Immuno-Blot PVDF membranes by wet transfer, and membranes were blocked in 5% (w/v) milk in TBS-T for 1 hour at room temperature. Primary antibodies were added and incubated at 4 °C overnight, followed by washing in TBS-T, incubation in secondary antibody for 1 hour at room temperature, and washing in TBS-T. Chemiluminescence detection of HRP-conjugated secondary antibody was performed using a cECL Western Blot Kit (CWBIO, CW0048). The images were acquired by Amersham Imager 680.

### Fluorescence microscopy

For live-cell microscopy, cells were replated into glass-bottom dishes (Beyotime, FCFC020) after siRNA transfection and transient plasmid transfection using the same procedure as for western blotting. The cells were imaged on a Zeiss LSM900 confocal microscope with a Plan-Apochromat 63x/1.40 oil objective.

For the colocalization assay, HeLa cells stably expressing Flag-GFP-TMEM192 were grown onto 20-mm glass coverslips for transient transfection the next day. Twenty-four hours after transfection, the cells were fixed with 4% v/v paraformaldehyde in PBS for 20 minutes and permeabilized with 0.1% (v/v) Triton X-100 for 20 minutes. Slides were mounted in anti-fade mounting medium with DAPI (Beyotime, P0131). Digital images were captured with a Zeiss LSM900 confocal microscope with a Plan-Apochromat 63x/1.40 oil objective. The term colocalization refers to the coincident detection of above-background green and red fluorescent signals in the same region.

### Data analysis and statistics

Late endosome/lysosome diameters were measured using ZEN (Blue edition) software with late endosomes/lysosomes approximated as circles. The selection of late endosomes/lysosomes was random. Clustered late endosomes/lysosomes were not included in the size analysis. Data were analyzed using GraphPad 9.0.0 software. Differences between datasets were analyzed using two-way analysis of variance (ANOVA), and P values of < 0.05 were considered statistically significant. *: P < 0.05; **: P < 0.01; ***: P<0.001; ****: P < 0.0001.

---

**Figure S1:**
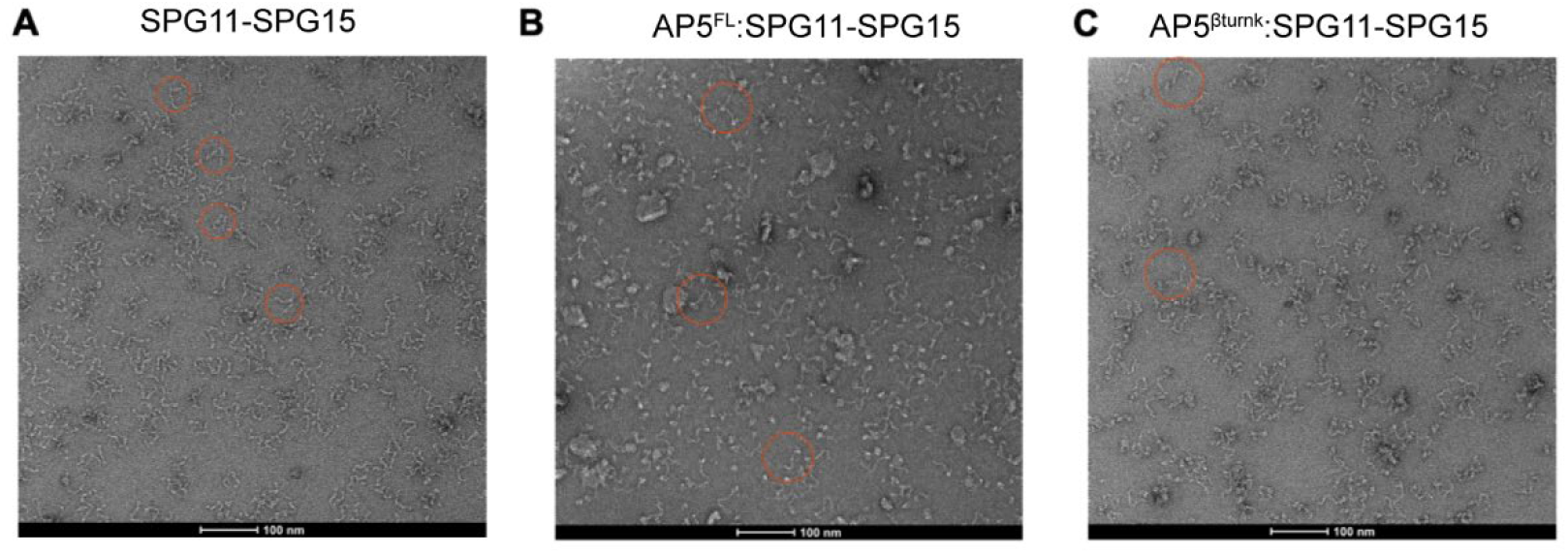
Representative negative stain micrographs of SPG11-SPG15, AP5^FL^:SPG11-SPG15 and AP5^βtrunk^:SPG11-SPG15 complex. The shape of the complex can be recognized and circled in the raw image.

**Figure S2:**
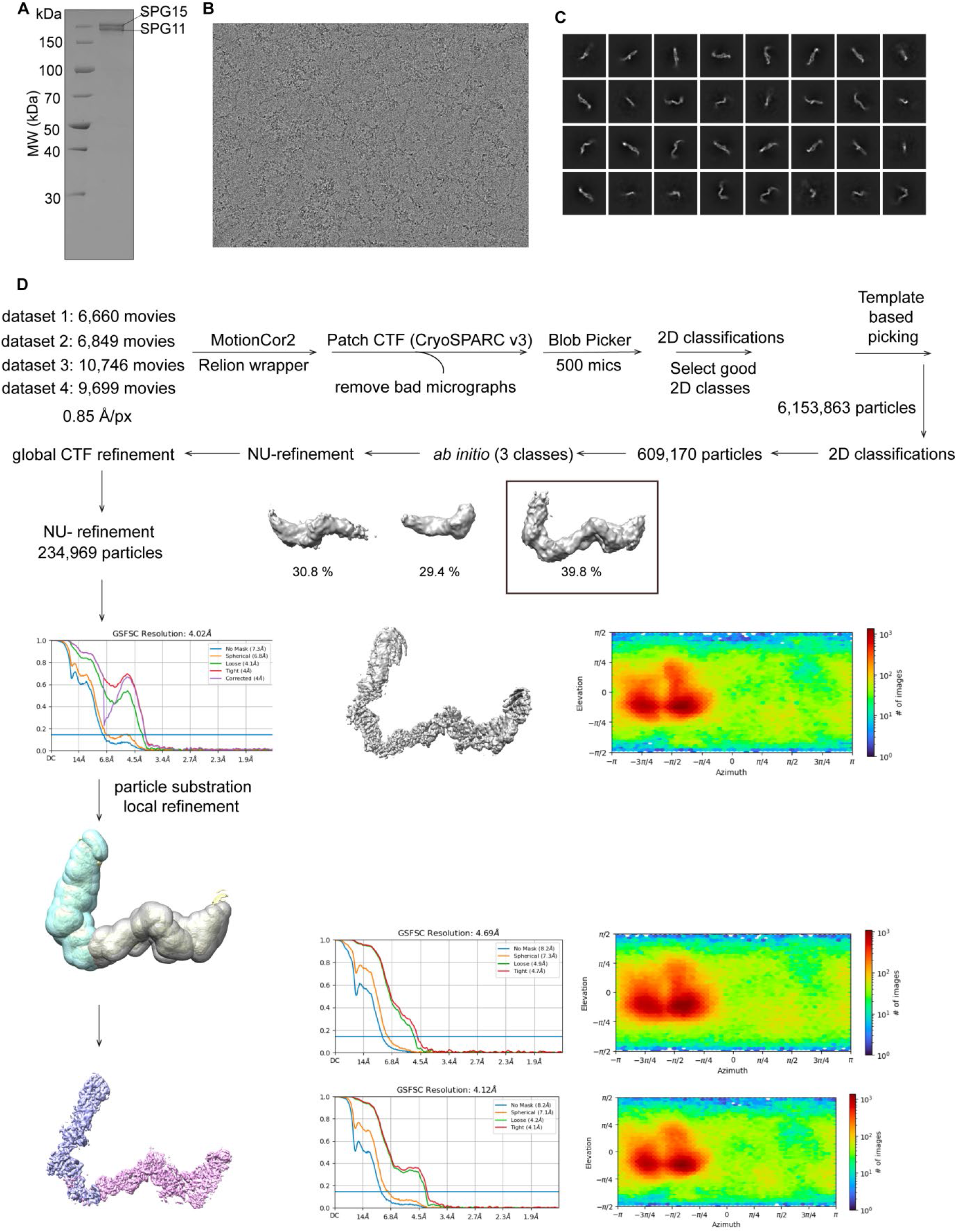
Overview of cryo-EM processing of human SPG11-SPG15. **(A)** Coomassie blue-stained SDS-PAGE of purified SPG11-SPG15. MW, molecular weight. **(B)** Representative motion-corrected cryo-EM micrograph of the human SPG11-SPG15 complex. **(C)** Representative 2D class averages for the SPG11-SPG15 complex. **(D)** Flow chart of cryo-EM data processing.

**Figure S3:**
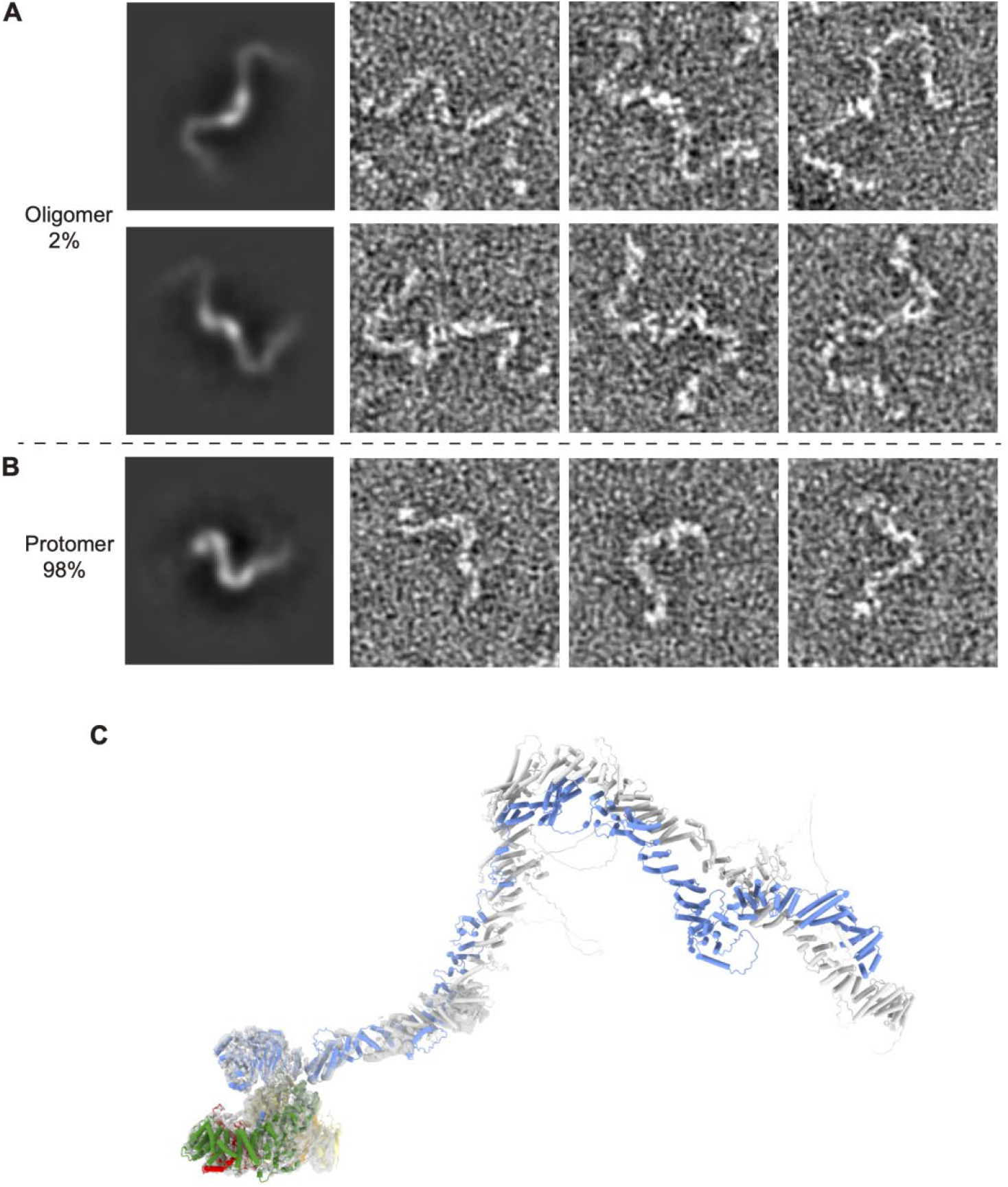
Negative stain analysis of SPG11-SPG15. **(A-B)** The 2D class averages and representative particles showing the dimeric SPG11-SPG15 and monomer SPG11-SPG15. **(C)** The model of the AP5^FL^:SPG11-SPG15 complex fitted into the map.

**Figure S4:**
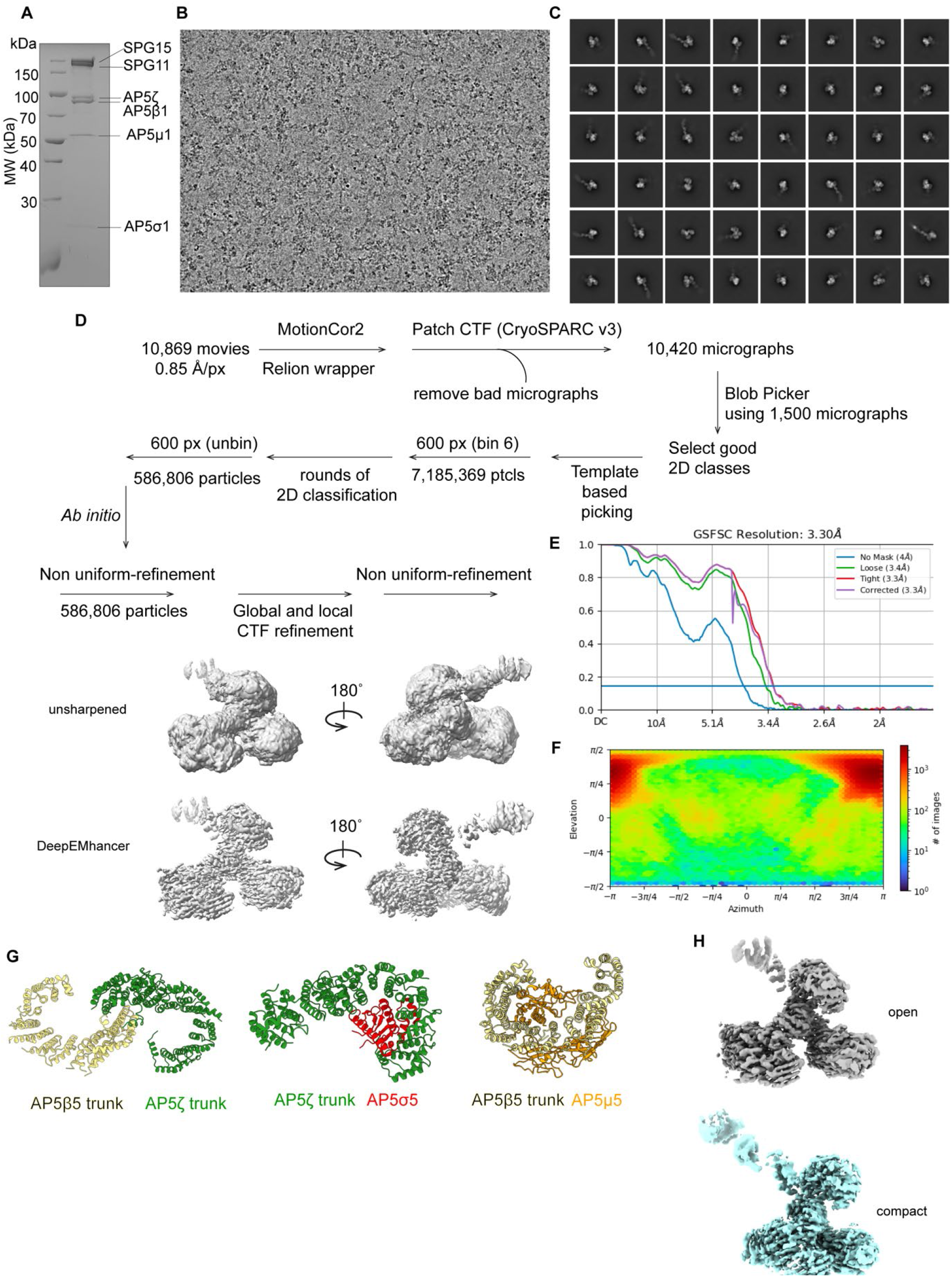
Cryo-EM structure determination of the AP5:SPG11-SPG15. **(A)** Coomassie blue-stained SDS-PAGE of the purified AP5:SPG11-SPG15 complex. MW, molecular weight. **(B)** Representative motion-corrected cryo-EM micrograph of the AP5:SPG11-SPG15 complex. **(C)** Representative 2D class averages for the AP5:SPG11-SPG15 complex. **(D)** Flow chart of cryo-EM data processing. NU-refinement: nonuniform refinement. **(E)** The FSC plots are between two independently refined half-maps with no mask (blue), spherical mask (orange), loose mask (green), tight mask (red), and corrected (purple). A cut-off of 0.143 (blue line) was used to estimate the resolution. **(F)** Angular particle distribution calculated in cryoSPARC for particle projections. The heatmap shows the number of particles for each viewing angle. **(G)** The structure of the AP5β5 trunk in complex with the ζ trunk, the ζ trunk in complex with σ5 and the β5 trunk in complex with μ5. **(H)** Open and compact conformation of the AP5 complex.

**Figure S5:**
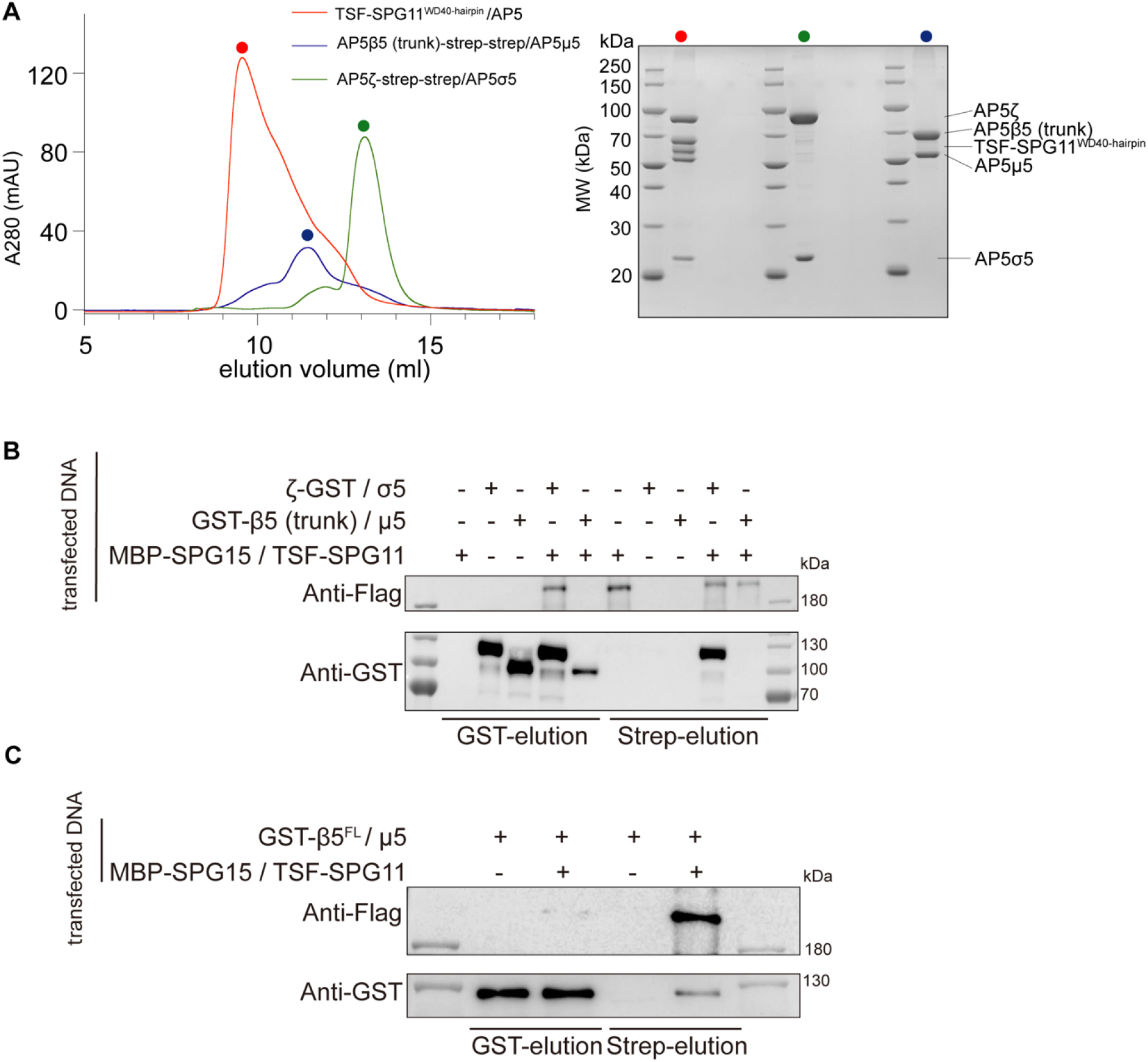
SPG11 is required for AP5 complex assembly. **(A)** Size exclusion profiles (Superdex 200 Increased 10/300 GL) of AP5ζ/σ5, AP5β5^trunk^/µ5 subcomplexes as well as incubation with SPG11^WD40-hairpin^ (*left*). SDS-PAGE analysis of the peak fractions from the complexes (*right*). **(B)** Pull-down experiment of MBP-SPG15, TSF-SPG11, GST-AP5β5^trunk^/µ5 and AP5ζ-GST/σ5. **(C)** Pull-down experiment of MBP-SPG15, TSF-SPG11 and GST-AP5β5/µ5.

**Figure S6:**
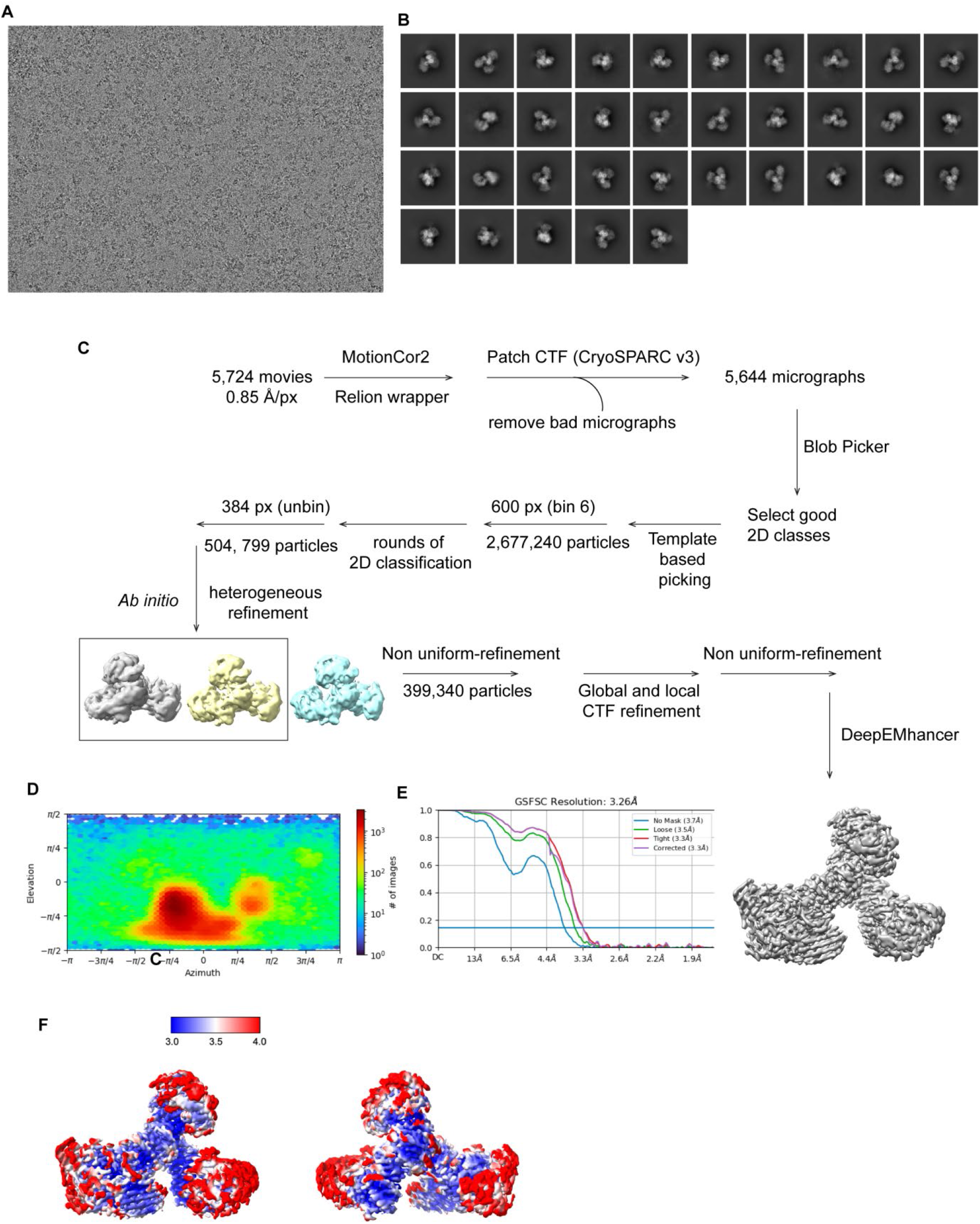
Cryo-EM structure determination of the AP5^βtrunk^:SPG11-SPG15 complex. **(A)** Representative motion-corrected cryo-EM micrograph of the AP5^βtrunk^:SPG11-SPG15 complex. **(B)** Representative 2D class averages for the AP5^βtrunk^:SPG11-SPG15 complex. **(C)** Flow chart of cryo-EM data processing. NU-refinement: nonuniform refinement. **(D)** Angular particle distribution calculated in cryoSPARC for particle projections. The heatmap shows the number of particles for each viewing angle. **(E)** The FSC plots are between two independently refined half-maps with no mask (blue), spherical mask (orange), loose mask (green), tight mask (red), and corrected (purple). A cut-off of 0.143 (blue line) was used to estimate the resolution. **(F)** AP5^βtrunk^:SPG11-SPG15 map color-coded by the local resolution estimation.

**Figure S7:**
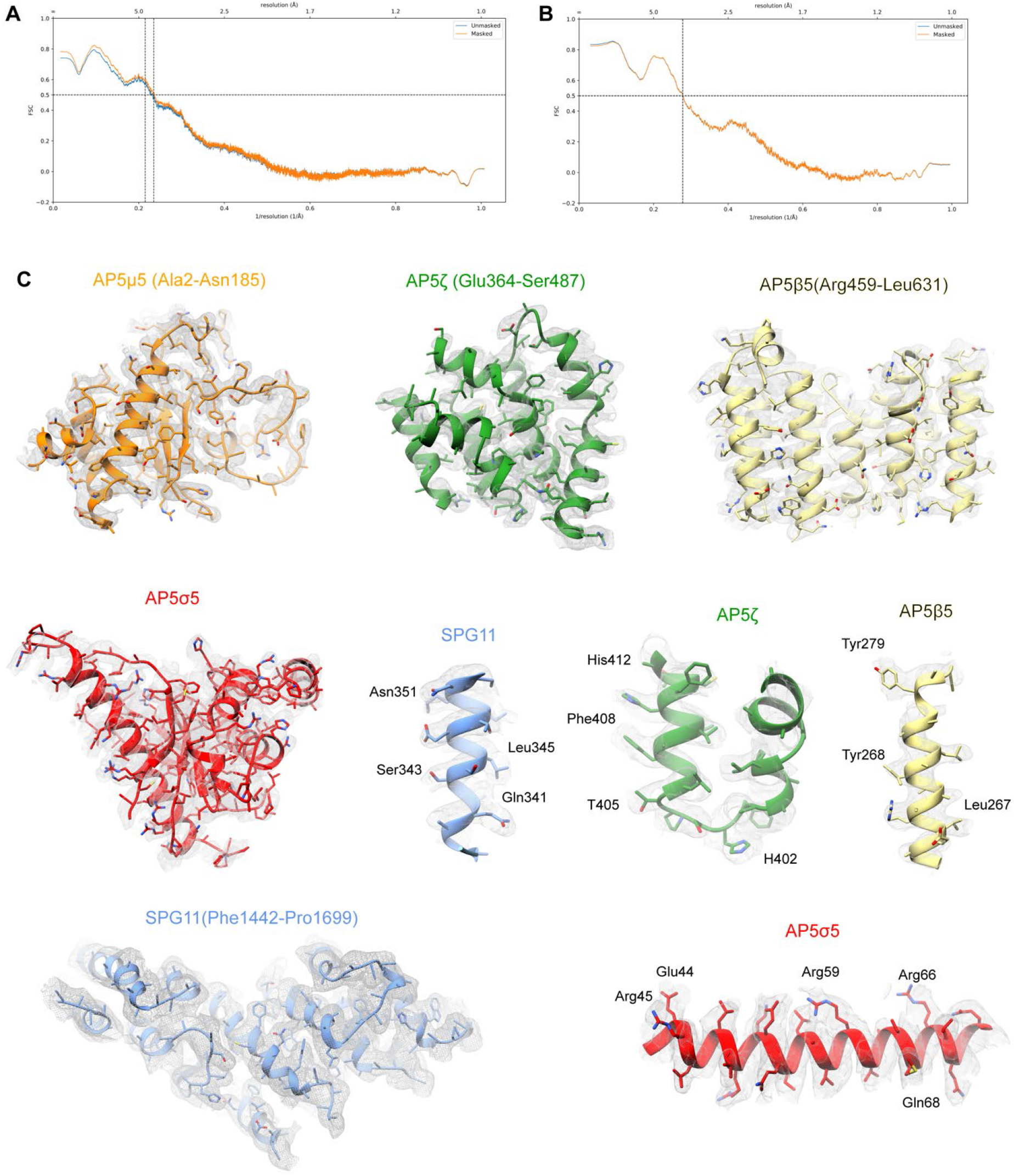
Model to map fitting. **(A)** FSC between the model and map for SPG11-SPG15 against the cryo-EM map. **(B)** FSC between the model and map for the AP5^βtrunk^:SPG11-SPG15 complex against the cryo-EM map. **(C)** Representative cryo-EM densities fitted to the model.

**Figure S8:**
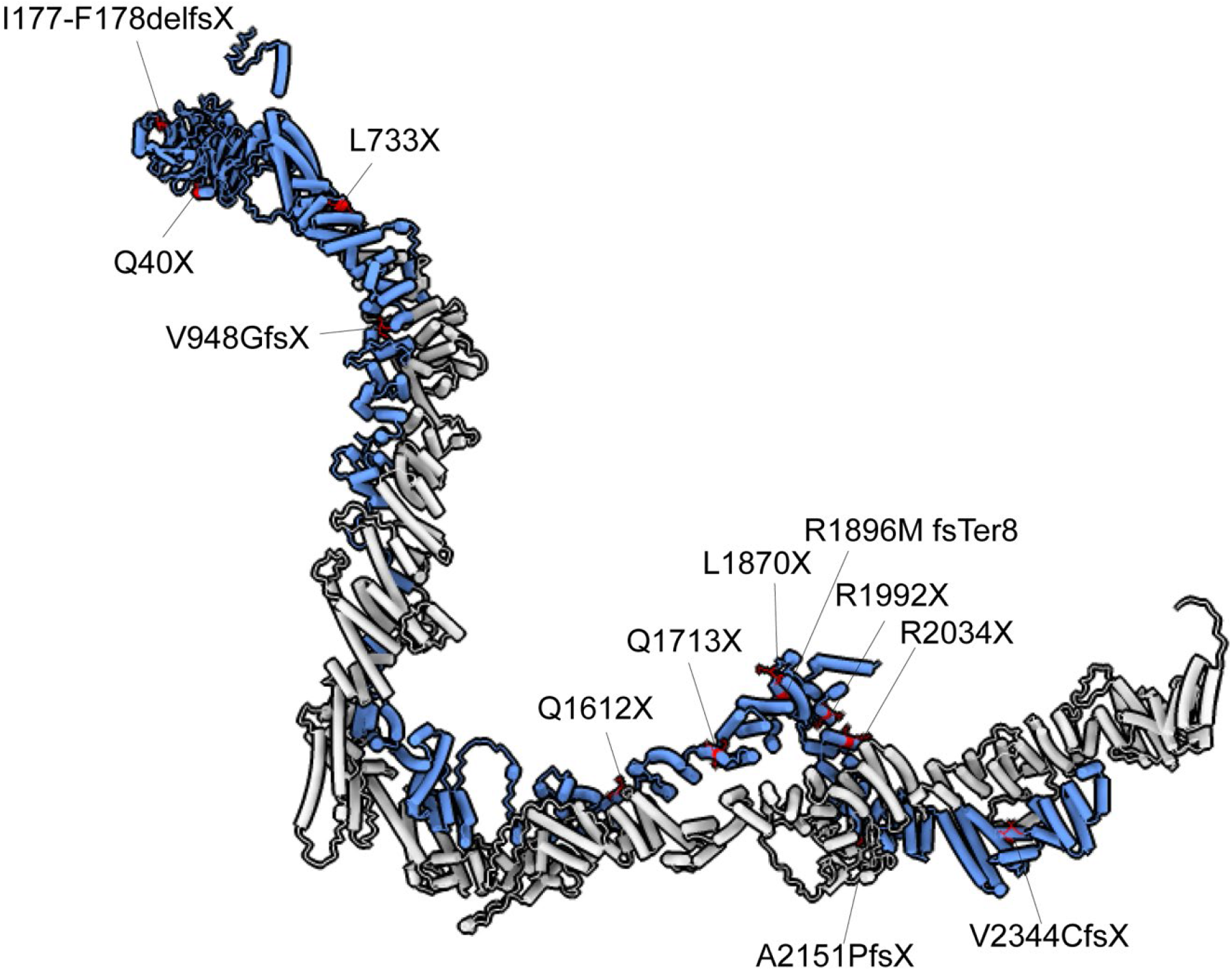
SPG11 mutations associated with spastic paraplegia mapped onto the SPG11-SPG15 structure. Many of the SPG11 mutations causative of spastic paraplegia are predicted to result in premature termination of peptide synthesis [2, 40, 41].

**Figure S9:**
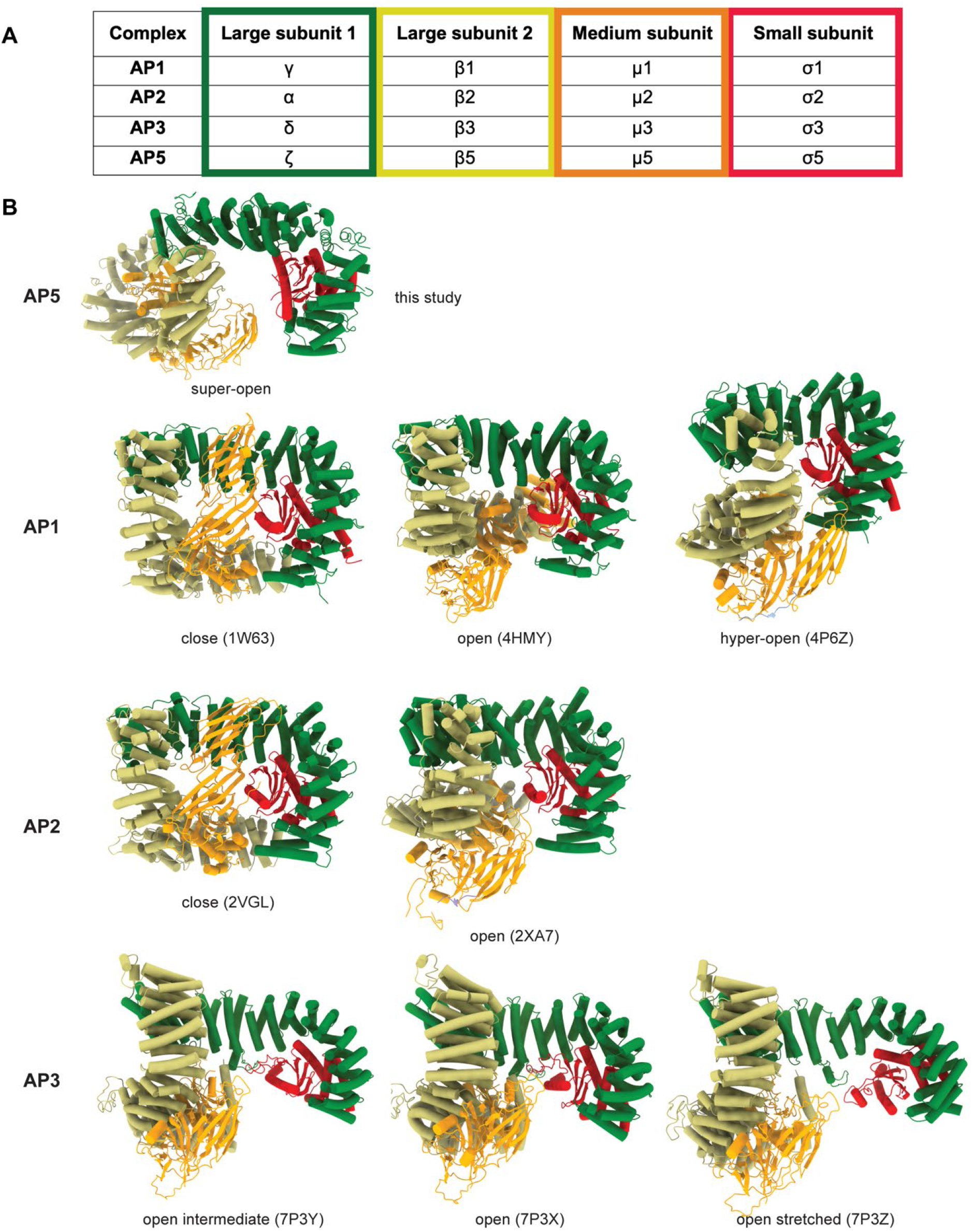
Structural comparison of the AP5 complex with AP1-3 complexes. **(A)** List of subunits in the four AP complexes including AP1-3 and AP5 complex. **(B)** Conformational states for AP1-3 and AP5 complexes.

**Figure S10:**
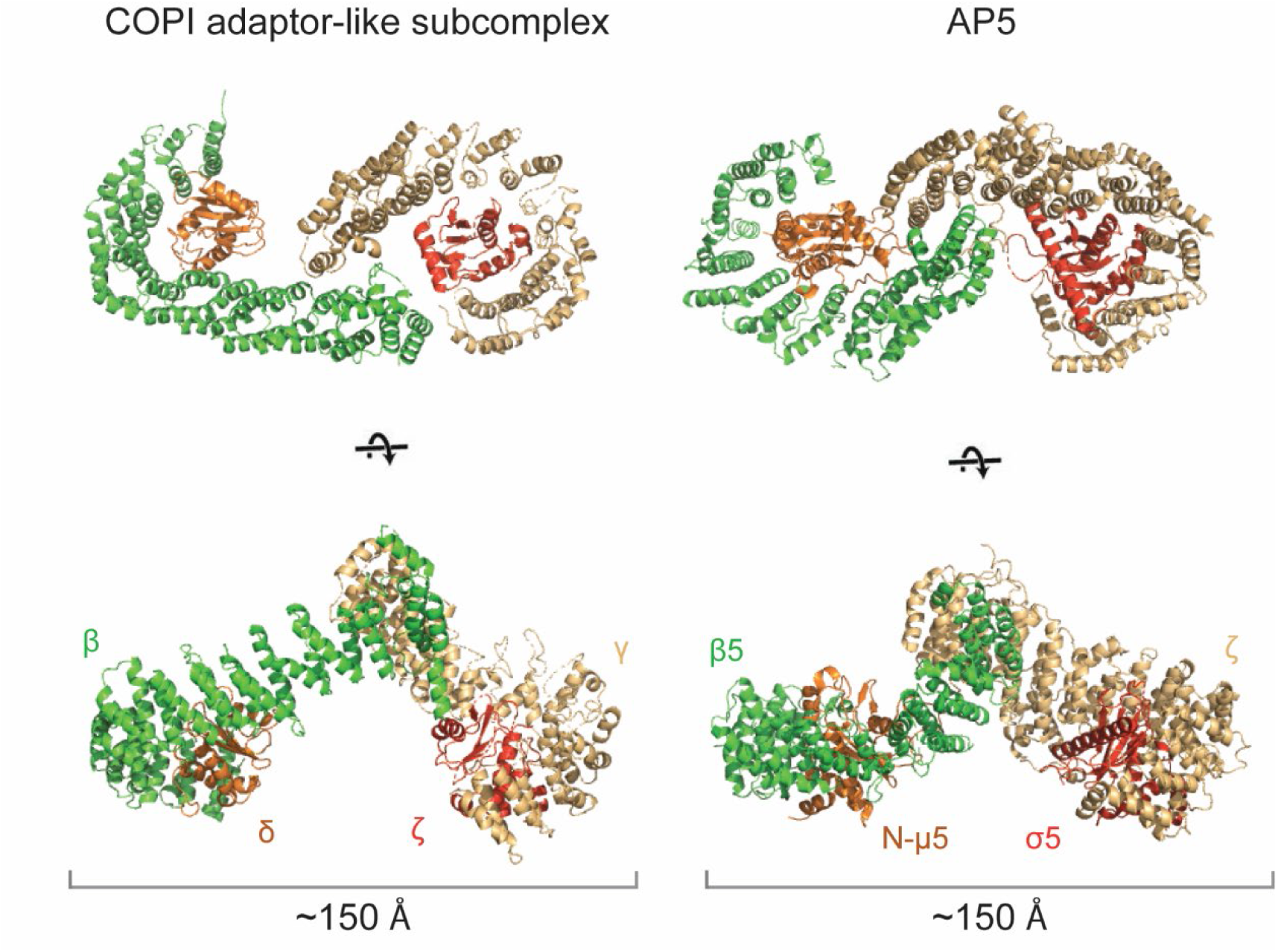
Structural comparison of the AP5 complex with the adaptor-like γ-ζ-β-δ-COP subcomplex of COPI complex. AP5 is in super-open conformation with two arms of the complex measuring ∼150Å apart. Structure model of the extended γ-ζ-β-δ-COP subcomplex is shown for comparison (PDB ID 5NZT). The ear domains of γ-COP and β-COP as well as C-µ5 were removed for clarity.

**Figure S11:**
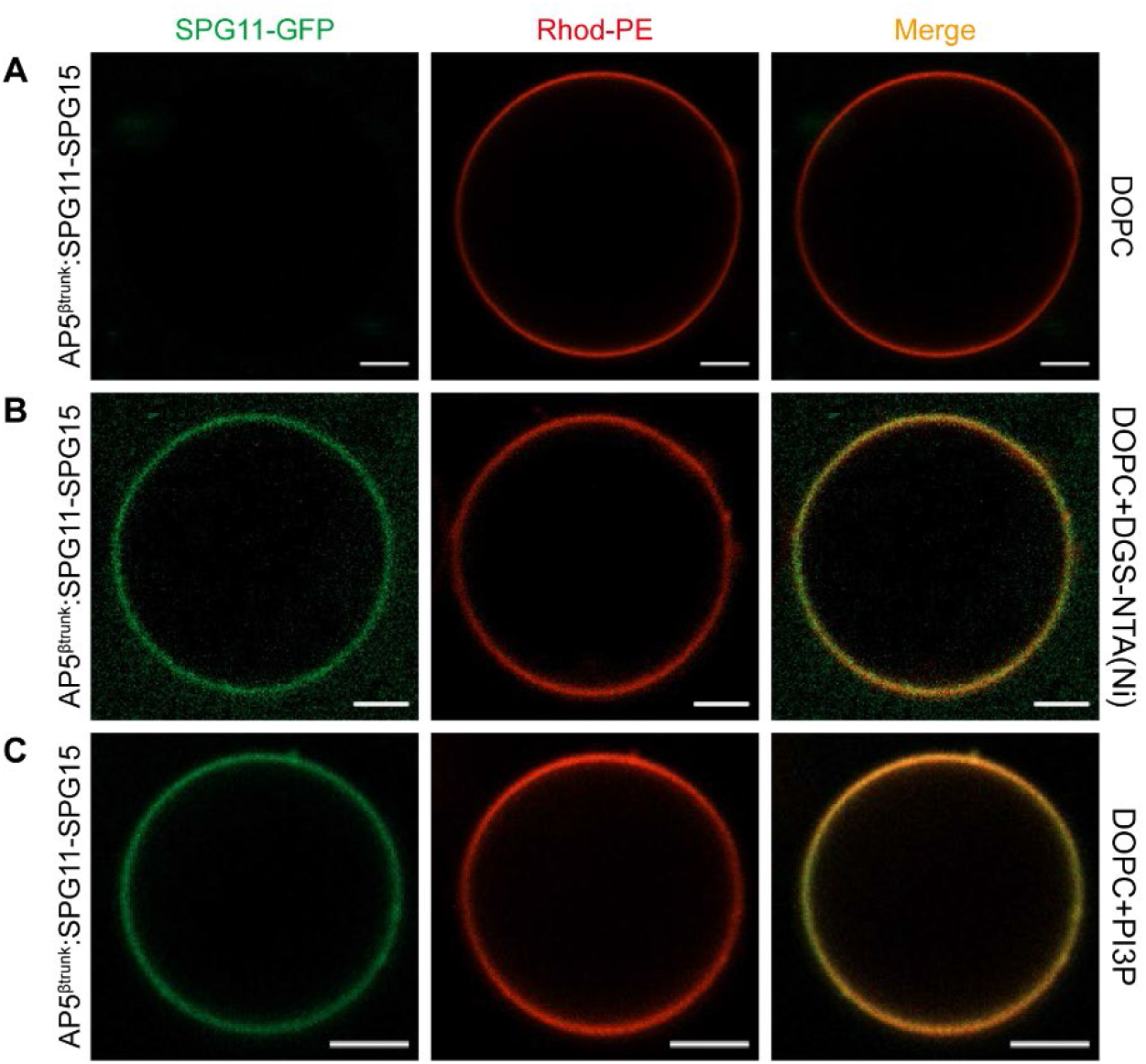
Effect of AP5^βtrunk^:SPG11-SPG15 complex on GUVs with different lipid composition. **(A)** AP5^βtrunk^:SPG11-SPG15 complex do not bind GUVs without PI3P and DGS-NTA(Ni). **(B), (C)** AP5^βtrunk^:SPG11-SPG15 complex can bind to GUVs with PI3P or DGS-NTA(Ni) but did not remodel GUV membrane. Each experiment repeated independently 3 times with similar results. All scale bars are 5 μm.

## References

1. Meyyazhagan, A., H. Kuchi Bhotla, M. Pappuswamy, and A. Orlacchio, The Puzzle of Hereditary Spastic Paraplegia: From Epidemiology to Treatment. Int J Mol Sci, 2022. 23(14).

2. Stevanin, G., et al., Mutations in SPG11, encoding spatacsin, are a major cause of spastic paraplegia with thin corpus callosum. Nat Genet, 2007. 39(3): p. 366–72.

3. Stevanin, G., et al., Mutations in SPG11 are frequent in autosomal recessive spastic paraplegia with thin corpus callosum, cognitive decline and lower motor neuron degeneration. Brain, 2008. 131(Pt 3): p. 772–84.

4. Goizet, C., et al., SPG15 is the second most common cause of hereditary spastic paraplegia with thin corpus callosum. Neurology, 2009. 73(14): p. 1111–1119.

5. Stenson, P.D., et al., Human Gene Mutation Database (HGMD): 2003 update. Hum Mutat, 2003. 21(6): p. 577–81.

6. Hirst, J., et al., Role of the AP-5 adaptor protein complex in late endosome-to-Golgi retrieval. PLoS Biol, 2018. 16(1): p. e2004411.

7. Khundadze, M., et al., A mouse model for SPG48 reveals a block of autophagic flux upon disruption of adaptor protein complex five. Neurobiology of Disease, 2019. 127: p. 419–431.

8. Hirst, J., et al., Interaction between AP-5 and the hereditary spastic paraplegia proteins SPG11 and SPG15. Mol Biol Cell, 2013. 24(16): p. 2558–69.

9. Murmu, R.P., et al., Cellular distribution and subcellular localization of spatacsin and spastizin, two proteins involved in hereditary spastic paraplegia. Mol Cell Neurosci, 2011. 47(3): p. 191–202.

10. Vantaggiato, C., et al., Rescue of lysosomal function as therapeutic strategy for SPG15 hereditary spastic paraplegia. Brain, 2023. 146(3): p. 1103–1120.

11. Edmison, D., L. Wang, and S. Gowrishankar, Lysosome Function and Dysfunction in Hereditary Spastic Paraplegias. Brain Sci, 2021. 11(2).

12. Varga, R.E., et al., In Vivo Evidence for Lysosome Depletion and Impaired Autophagic Clearance in Hereditary Spastic Paraplegia Type SPG11. PLoS Genet, 2015. 11(8): p. e1005454.

13. Renvoise, B., et al., Lysosomal abnormalities in hereditary spastic paraplegia types SPG15 and SPG11. Ann Clin Transl Neurol, 2014. 1(6): p. 379–389.

14. Chang, J., S. Lee, and C. Blackstone, Spastic paraplegia proteins spastizin and spatacsin mediate autophagic lysosome reformation. J Clin Invest, 2014. 124(12): p. 5249–62.

15. Hirst, J., et al., Loss of AP-5 results in accumulation of aberrant endolysosomes: defining a new type of lysosomal storage disease. Human Molecular Genetics, 2015. 24(17): p. 4984–4996.

16. Nixon, R.A., The role of autophagy in neurodegenerative disease. Nat Med, 2013. 19(8): p. 983–97.

17. Park, H., J.H. Kang, and S. Lee, Autophagy in Neurodegenerative Diseases: A Hunter for Aggregates. International Journal of Molecular Sciences, 2020. 21(9).

18. Menzies, F.M., et al., Autophagy and Neurodegeneration: Pathogenic Mechanisms and Therapeutic Opportunities. Neuron, 2017. 93(5): p. 1015–1034.

19. Malik, B.R., D.C. Maddison, G.A. Smith, and O.M. Peters, Autophagic and endo-lysosomal dysfunction in neurodegenerative disease. Molecular Brain, 2019. 12(1).

20. Hirst, J., et al., Interaction between AP-5 and the hereditary spastic paraplegia proteins SPG11 and SPG15. Molecular Biology of the Cell, 2013. 24(16): p. 2558–2569.

21. Hirst, J., G.G. Hesketh, A.C. Gingras, and M.S. Robinson, Rag GTPases and phosphatidylinositol 3-phosphate mediate recruitment of the AP-5/SPG11/SPG15 complex. J Cell Biol, 2021. 220(2).

22. Dacks, J.B. and M.S. Robinson, Outerwear through the ages: evolutionary cell biology of vesicle coats. Current Opinion in Cell Biology, 2017. 47: p. 108–116.

23. Tan, J.Z.A. and P.A. Gleeson, Cargo Sorting at the trans-Golgi Network for Shunting into Specific Transport Routes: Role of Arf Small G Proteins and Adaptor Complexes. Cells, 2019. 8(6).

24. Sanger, A., J. Hirst, A.K. Davies, and M.S. Robinson, Adaptor protein complexes and disease at a glance. Journal of Cell Science, 2019. 132(20).

25. Jackson, L.P., et al., A Large-Scale Conformational Change Couples Membrane Recruitment to Cargo Binding in the AP2 Clathrin Adaptor Complex. Cell, 2010. 141(7): p. 1220–U213.

26. Ren, X.F., et al., Structural Basis for Recruitment and Activation of the AP-1 Clathrin Adaptor Complex by Arf1. Cell, 2013. 152(4): p. 755–767.

27. Kelly, B.T., et al., AP2 controls clathrin polymerization with a membrane-activated switch. Science, 2014. 345(6195): p. 459–463.

28. Beacham, G.M., E.A. Partlow, and G. Hollopeter, Conformational regulation of AP1 and AP2 clathrin adaptor complexes. Traffic, 2019. 20(10): p. 741–751.

29. Punjani, A. and D.J. Fleet, 3D variability analysis: Resolving continuous flexibility and discrete heterogeneity from single particle cryo-EM. J Struct Biol, 2021. 213(2): p. 107702.

30. Hayakawa, A., et al., Structural basis for endosomal targeting by FYVE domains. Journal of Biological Chemistry, 2004. 279(7): p. 5958–5966.

31. Dumas, J.J., et al., Multivalent endosome targeting by homodimeric EEA1. Molecular Cell, 2001. 8(5): p. 947–958.

32. Misra, S. and J.H. Hurley, Crystal structure of a phosphatidylinositol 3-phosphate-specific membrane-targeting motif, the FYVE domain of Vps27p. Cell, 1999. 97(5): p. 657–666.

33. Stenmark, H., R. Aasland, and P.C. Driscoll, The phosphatidylinositol 3-phosphate-binding FYVE finger. Febs Letters, 2002. 513(1): p. 77–84.

34. Kutateladze, T. and M. Overduin, Structural mechanism of endosome docking by the FYVE domain. Science, 2001. 291(5509): p. 1793–1796.

35. Dodonova, S.O., et al., 9 angstrom structure of the COPI coat reveals that the Arf1 GTPase occupies two contrasting molecular environments. Elife, 2017. 6.

36. Fath, S., J.D. Mancias, X.P. Bi, and J. Goldberg, Structure and organization of coat proteins in the COPII cage. Cell, 2007. 129(7): p. 1325–1336.

37. ter Haar, E., A. Musacchio, S.C. Harrison, and T. Kirchhausen, Atomic structure of clathrin: A beta propeller terminal domain joins an alpha zigzag linker. Cell, 1998. 95(4): p. 563–573.

38. Renvoise, B., et al., Lysosomal abnormalities in hereditary spastic paraplegia types SPG15 and SPG11. Annals of Clinical and Translational Neurology, 2014. 1(6): p. 379–389.

39. Chang, J., S. Lee, and C.D. Blackstone, Spastic paraplegia proteins spastizin and spatacsin mediate autophagic lysosome reformation. Molecular Biology of the Cell, 2014. 25.

40. Li, C., et al., Mild cognitive impairment in novel SPG11 mutation-related sporadic hereditary spastic paraplegia with thin corpus callosum: case series. BMC Neurol, 2021. 21(1): p. 12.

41. Duan, J.Q., H. Liu, and J.Q. Wu, Case report: Novel mutations in the SPG11 gene in a case of autosomal recessive hereditary spastic paraplegia with a thin corpus callosum. Front Integr Neurosci, 2023. 17: p. 1117617.

42. Collins, B.M., et al., Molecular architecture and functional model of the endocytic AP2 complex. Cell, 2002. 109(4): p. 523–535.

43. Hirst, J., et al., Role of the AP-5 adaptor protein complex in late endosome-to-Golgi retrieval. Plos Biology, 2018. 16(1).

44. Taylor, R.J., G. Tagiltsev, and J.A.G. Briggs, The structure of COPI vesicles and regulation of vesicle turnover. Febs Letters, 2023. 597(6): p. 819–835.

45. Dodonova, S.O., et al., A structure of the COPI coat and the role of coat proteins in membrane vesicle assembly. Science, 2015. 349(6244): p. 195–198.

46. Harakuge, S., En-Bloc Incorporation of Coatomer Subunits during the Assembly of Cop- Coated Vesicles (Vol 124, Pg 883, 1994). Journal of Cell Biology, 1994. 126(2): p. 589–589.

47. Carlton, J.G. and P.J. Cullen, Coincidence detection in phosphoinositide signaling. Trends in Cell Biology, 2005. 15(10): p. 540–547.

48. Heldwein, E.E., et al., Crystal structure of the clathrin adaptor protein 1 core. Proceedings of the National Academy of Sciences of the United States of America, 2004. 101(39): p. 14108–14113.

49. Kadlecova, Z., et al., Regulation of clathrin-mediated endocytosis by hierarchical allosteric activation of AP2. Journal of Cell Biology, 2017. 216(1): p. 167–179.

50. Honing, S., et al., Phosphatidylinositol-(4,5)-bisphosphate regulates sorting signal recognition by the clathrin-associated adaptor complex AP2 (vol 19, pg 577, 2005). Molecular Cell, 2005. 19(4): p. 577–577.

51. Baust, T., et al., Protein networks supporting AP-3 function in targeting lysosomal membrane proteins. Molecular Biology of the Cell, 2008. 19(5): p. 1942–1951.

52. Schoppe, J., et al., Flexible open conformation of the AP-3 complex explains its role in cargo recruitment at the Golgi. Journal of Biological Chemistry, 2021. 297(5).

53. Boutry, M., et al., Loss of spatacsin impairs cholesterol trafficking and calcium homeostasis. Communications Biology, 2019. 2.

54. Wang, S., et al., Structural characterization of coatomer in its cytosolic state. Protein Cell, 2016. 7(8): p. 586–600.

55. Antonny, B., Membrane deformation by protein coats. Curr Opin Cell Biol, 2006. 18(4): p. 386–94.

56. Nanayakkara, R., et al., Autophagic lysosome reformation in health and disease. Autophagy, 2023. 19(5): p. 1378–1395.

57. Leneva, N., et al., Architecture and mechanism of metazoan retromer:SNX3 tubular coat assembly. Sci Adv, 2021. 7(13).

58. Chi, R.J., M.S. Harrison, and C.G. Burd, Biogenesis of endosome-derived transport carriers. Cell Mol Life Sci, 2015. 72(18): p. 3441–3455.

59. Buser, D.P. and A. Spang, Protein sorting from endosomes to the TGN. Front Cell Dev Biol, 2023. 11: p. 1140605.

60. Weeratunga, S., B. Paul, and B.M. Collins, Recognising the signals for endosomal trafficking. Curr Opin Cell Biol, 2020. 65: p. 17–27.

61. Vantaggiato, C., et al., ZFYVE26/SPASTIZIN and SPG11/SPATACSIN mutations in hereditary spastic paraplegia types AR-SPG15 and AR-SPG11 have different effects on autophagy and endocytosis. Autophagy, 2019. 15(1): p. 34–57.

62. Punjani, A., J.L. Rubinstein, D.J. Fleet, and M.A. Brubaker, cryoSPARC: algorithms for rapid unsupervised cryo-EM structure determination. Nat Methods, 2017. 14(3): p. 290–296.

63. Zheng, S.Q., et al., MotionCor2: anisotropic correction of beam-induced motion for improved cryo-electron microscopy. Nat Methods, 2017. 14(4): p. 331–332.

64. Zivanov, J., et al., New tools for automated high-resolution cryo-EM structure determination in RELION-3. Elife, 2018. 7.

65. Sanchez-Garcia, R., et al., DeepEMhancer: a deep learning solution for cryo-EM volume post-processing. Commun Biol, 2021. 4(1): p. 874.

66. Jumper, J., et al., Highly accurate protein structure prediction with AlphaFold. Nature, 2021. 596(7873): p. 583–589.

67. Emsley, P. and K. Cowtan, Coot: model-building tools for molecular graphics. Acta Crystallogr D Biol Crystallogr, 2004. 60(Pt 12 Pt 1): p. 2126-32.

68. Pettersen, E.F., et al., UCSF Chimera--a visualization system for exploratory research and analysis. J Comput Chem, 2004. 25(13): p. 1605–12.

69. Adams, P.D., et al., PHENIX: a comprehensive Python-based system for macromolecular structure solution. Acta Crystallogr D Biol Crystallogr, 2010. 66(Pt 2): p. 213–21.

70. Afonine, P.V., et al., Real-space refinement in PHENIX for cryo-EM and crystallography. Acta Crystallogr D Struct Biol, 2018. 74(Pt 6): p. 531–544.

71. Chen, V.B., et al., MolProbity: all-atom structure validation for macromolecular crystallography. Acta Crystallogr D Biol Crystallogr, 2010. 66(Pt 1): p. 12–21.

72. Pettersen, E.F., et al., UCSF ChimeraX: Structure visualization for researchers, educators, and developers. Protein Sci, 2021. 30(1): p. 70–82.

